# Role of the unstructured N-linker of the alkaline phosphatase superfamily member BcsG in the gastrointestinal pathogen *Salmonella typhimurium*

**DOI:** 10.64898/2026.09.23.753746

**Authors:** Li Li, Ute Römling

## Abstract

A subgroup of membrane-anchored bacterial Alkaline Phosphatase superfamily members transfers phospholipid headgroups from phospholipids to diverse acceptor molecules. As part of type II and hybrid type I/II cellulose biosynthesis gene clusters BcsG transfers phosphorylethanolamine pEtN from phospholipid to the emerging 1,4 β-D-glucan chain synthesized by the BcsABC cellulose biosynthesis nanomachine. BcsG homologs throughout the phylogenetic tree possess a uniquely structured tripartite linker that connects the fifth transmembrane domain with the periplasmic catalytic domain. A N-terminal unstructured sequence of highly variable length and biased amino acid composition, the N-linker, is followed by an α-helix and an unstructured sequence of high similarity and constant length leading into the first β-strand of the catalytic domain. With BcsG from *Salmonella typhimurium* synthesizing pEtN cellulose as a model, linkers were shown to be promiscuous and linker length positively correlated with transfer efficiency of the pEtN headgroup and protein stability. As the donor substrate, the phosphatidylethanolamine content of the membranes affects pEtN transfer efficiency. With Alkaline Phosphatase superfamily members to be promiscuous enzymes the catalytic domain of OpgE transfers pEtN to osmoregulated periplasmic glucan. BcsG-OpgE hybrid proteins cause colony morphology alterations thus potentially transferring pEtN to the BcsA synthesized glucan chain. With BcsG with shortest linkers not associated with a cellulose biosynthesis gene cluster, multiple hypotheses can be proposed that select for different length of the N-linker in BcsG homologs located within a cellulose biosynthesis gene cluster in other Pseudomonadati.

**Importance:** Bacterial cellulose biosynthesis is performed by the cellulose synthase as the core catalytic entity with additional accessory gene product dependent on the type of organism. A number of Gram-negative organisms synthesize, however, covalently modified glucan chains by the action of enzymes co-localized with the cellulose synthase via their membrane part, while their catalytic domain is located in the periplasm. The role of unstructured linkers in proteins remains understudied. This work shows the impact of the unstructured linker region connecting the membrane part with the catalytic domain on apparent catalytic activity and protein level. Linker length is positively correlated with protein steady state levels and negatively with apparent transfer activity/cellulose biosynthesis.

## Introduction

The superfamily of alkaline phosphatase domains involved in phosphate and sulfate metabolism occurs in all domains of life including bacteria, archaea and, in eukaryotes, fungi, algae and animals, but is largely absent in higher land plants. With an overall α/β/α sandwich structure of the domain the catalytic site is located close to the surface of the protein which enables promiscuous catalytic activities (1, 2). The basic enzymatic activities of the superfamily members are phosphate/sulphate and phosphonate/sulphonate transferase and esterase activities with an individual enzyme to possess the ability to hydrolyze and eventually donate to structurally different substrates (3, 4). Despite substantial low sequence similarity, catalytic activity conventionally requires divalent cations such as Zn^2+^, Mg^2+^ and Mn^2+^ to form a metallic core at the catalytic site involved in positioning of the substrate and activation of the nucleophile. Besides the nucleophile which can be a serine or threonine residue and transition state stabilizing amino acids such as an arginine, other highly conserved amino acids such as histidines, aspartates and asparagines can further make up the catalytic site in different subgroups of the superfamily to contribute to substrate binding (1).

In Gram-negative bacteria, alkaline phosphatase superfamily members are found in the periplasmic space. While the prototype, the Alkaline Phosphatase encoded by *E. coli*, is a soluble protein, many alkaline phosphatase superfamily members are actually anchored in the cytoplasmic membrane connected with the catalytic domain by a long, partially unstructured and often tripartite linker sequence.

The gastrointestinal pathogen *Salmonella enterica* serovar Typhimurium codes for at least seven membrane anchored alkaline phosphatase superfamily members which possess low amino acid similarity (Fig. 1A). Three of these proteins, EptA, EptB and CptA and MCR, an occasionally horizontally transferred accessory protein mediating resistance towards the lipopeptide antibiotic colistin, are representatives of a subfamily dedicated to transfer phosphorylethanolamine phospholipid headgroups from phospholipid to the core of the lipopolysaccharide (LPS) to protect against membrane attacking polymyxins and antimicrobial peptides (5–12). Two enzymes, OpgB and OpgE, decorate the osmoregulated periplasmic glucan (OPG) with phosphorylglycerol and phosphorylethanolamine, respectively (13, 14). Of note, OpgB can be proteolytically processed and redistributes in its soluble periplasmic form the phosphoglycerol moiety among the glucose residues of the OPGs. PbgA actually has been reported to possess catalytic activity, but the protein mainly traffics cardiolipin glycerophospholipids from the inner to the outer membrane (7, 15). Last, but not least, BcsG decorates the emerging 1,4-β-D-glucan chain synthesized by the cellulose synthase BcsA on the OH group of the C6 carbon with phosphorylethanolamine (Fig. 1A; (2, 16)). The catalytic site of BcsG contains a pentavalent Zn^2+^ ion at the conserved Zn^2+^ binding site and uses a conserved serine 278 as the nucleophile.

**Fig 1.**
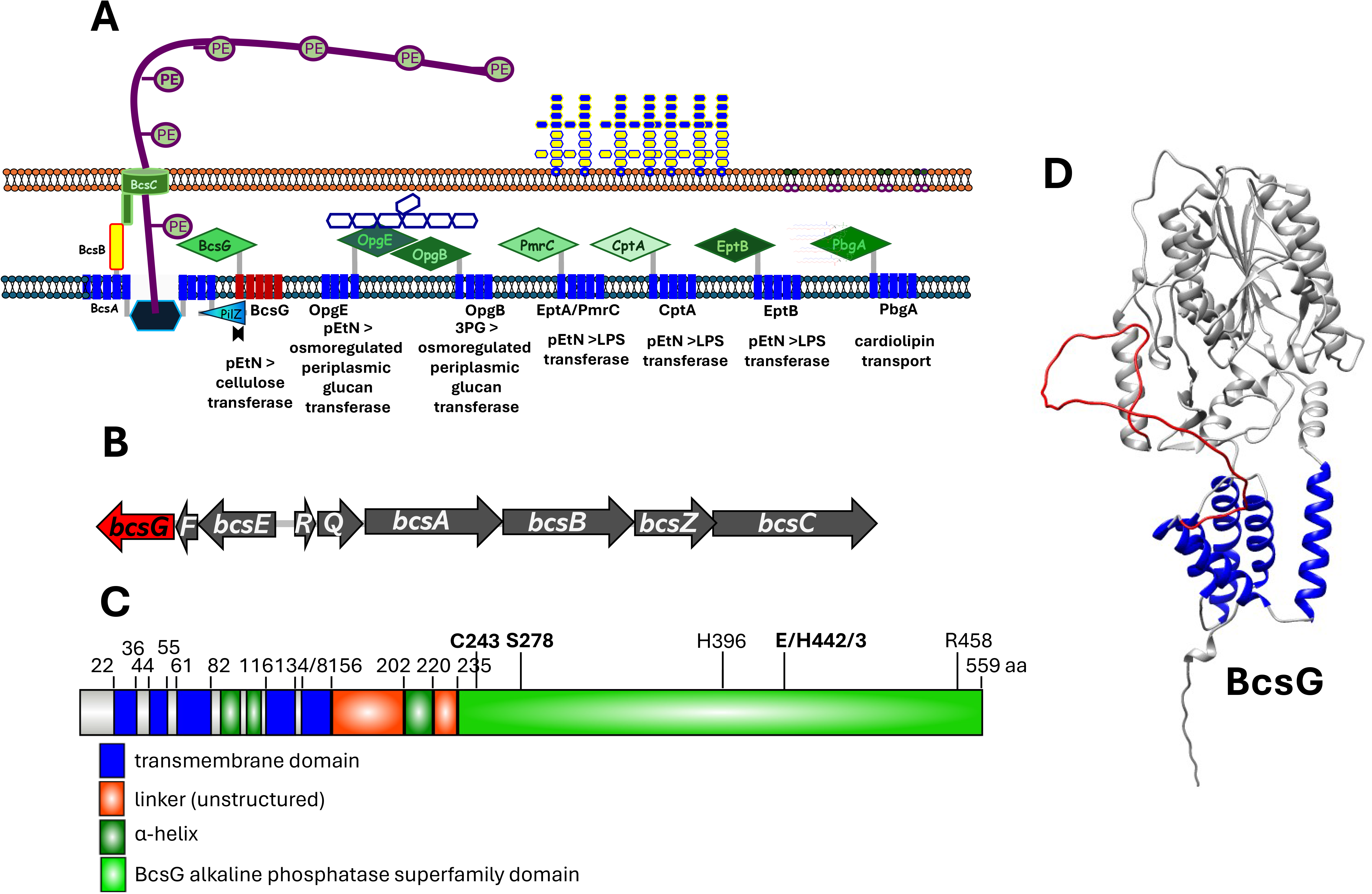
Alkaline phosphatase superfamily members of *S. typhimurium* ATCC 14028 and gene localization, domain structure and Alpha-Fold 3 model of BcsG. (A) Display of all seven alkaline phosphatase superfamily members of *S. typhimurium* ATCC 14028, OpgB, OpgE, EptA, EptB, CtpA, PbgA and BcsG. The accessory Mcr alkaline phosphatase superfamily member homologous to EptA, EptB and CtpA mediating colistin resistance is not displayed. (B) The type IIa cellulose biosynthesis gene cluster consists of two divergently transcribed operons *bcsQRABZC* and *bcsEFG*. (**C**) Domain composition and secondary structure elements outside of the catalytic domain of BcsG. (**D**) AlphaFold 3 model of BcsG. In dark blue, transmembrane helix; in red, N-linker connecting the 5^th^ transmembrane helix with the α-helix of the linker.

Cellulose is an exopolysaccharide produced as an extracellular matrix component upon biofilm formation by phylogenetically diverse bacteria (17–19). In Gram-negative bacteria a functional cellulose synthase is constituted by BcsA and BcsB, the catalytic subunit and the predominantly periplasmic BcsA-interacting component, respectively, as core components. Furthermore, cellulose biosynthesis modules contain a variety of variable accessory genetic elements that alter the chemical structure, conformation and higher order structure of the glucan chains. In this context, type II cellulose biosynthesis gene clusters contain a variably arranged *bcsEFG* module (Fig. 1B, (17)). BcsG has dual functionality in the type IIa cellulose biosynthesis gene cluster of *S. typhimurium* as its membrane part stabilizes the cellulose synthase BcsA (2).

Comparison of BcsG proteins from phylogenetically different bacteria indicted that the first third of the linker sequence connecting the fifth transmembrane helix with the catalytic domain has low sequence similarity, is highly variable in length and is structurally highly flexible, while the other two thirds of the linker sequence readily align with higher conservation of the amino acid sequence. We therefore started to investigate the phylogenetic and functional constraints that determine linker length by analysis of linker length distribution among BcsG homologs and by experimental assessment of the consequences of different linker lengths on phosphorylethanolamine decoration of cellulose by BcsG in *S. typhimurium*. As our results demonstrate that BcsG linker length is highly flexible without loosing its basic functionality, various hypothesis can explain the development of different linker length for BcsG homologs in phylogenetically diverse bacteria.

## RESULTS

### Structural modelling shows that BcsG has a tripartite periplasmic linker

*S. typhimurium* possesses seven highly diverse alkaline phosphatase superfamily members OpgB, OpgE, EptA, EptB, CptA, PbgA and BcsG which carry out distinct transfer reactions from phospholipid headgroups to various acceptor molecules (Fig. 1A; (5, 13, 14, 16)). Among the proteins that transfer the phosphorylethanolamine phospholipid headgroup, EptA, EptB and CptA transfer to the lipid A part of the lipopolysaccharide and OpgE transfers to 2’ C of osmoinduced periplasmic glucan. BcsG uniquely covalently links pEtN to glucose moieties of the glucan chain with a phosphodiester bond at the -OH group of the C6 position. Alphafold 3 modelling shows that all seven alkaline phosphatase superfamily members are predicted to possess four or five transmembrane helices at the N-terminal end, followed by a tripartite linker region and the C-terminal catalytic alkaline phosphatase domain with α/β/α sandwich organization (Fig. 1C and D; Fig. S1). The organization of the secondary structure of the linker region differs between the individual enzymes. While the first part of the linker sequence is highly variable, common is an approx. 16 amino acid long unstructured region at the 3’-end leading into the first β strand of the catalytic domain preceeded by an α-helix of variable length (Fig. 1C and D; Fig. S1). Of note, PbgA which transfers cardiolipin from the inner to the outer membrane is predicted to possess a predominantly unstructured linker sequence with a short α-helical part (Fig. 1).

In contrast to most other alkaline phosphatase superfamily members of *S. typhimurium*, the linker of BcsG has a clearly distinguishable tripartite structure with a 47 amino acid long unstructured sequence spanning from amino acid 156 to 202 (designated N-linker), an α-helical part from amino acid 203 to 220 and again an unstructured (guiding) sequence until amino acid 235 leading into the first β-strand of the central seven-stranded mixed β-sheet of the catalytic domain (Fig. 1C and D; (2)).

### The N-linker of BcsG is highly variable in length and amino acid composition

Upon investigating the first 5000 proteins homologous to BcsG of *S. typhimurium* retrieved by BLAST search and including the BcsG representative of type IIa-d cellulose biosynthesis gene clusters (17), we noticed, the 5’ located first third of the linker, the N-linker sequence, was highly variable in amino acid sequence and length (Fig. 2A-C; Fig. S2A) spanning from - 1 (according to BcsG_St_ linker definition) to 113 amino acids. However, the amino acid composition is highly biased with six amino acids, alanine and valine as amino acid with aliphatic side chains, glycine and proline as helix-breaking amino acids and serine and threonine as amino aids with polar side chains dominating with >70% of the total amino acids (Fig. S2B). The borders of the N-linker are clearly recognizable in all BcsG homologs as the amino acids at the N-terminus of the following α-helix contain highly conserved amino acid motifs (Fig. 2A; Fig. S2A). Otherwise, the linker is composed of semiconserved amino acid sequences in the 3’-located two-thirds of the C-terminal linker encompassing the α-helix and the unstructured guiding sequence leading into the catalytic domain (Fig. S2D).

**Fig 2.**
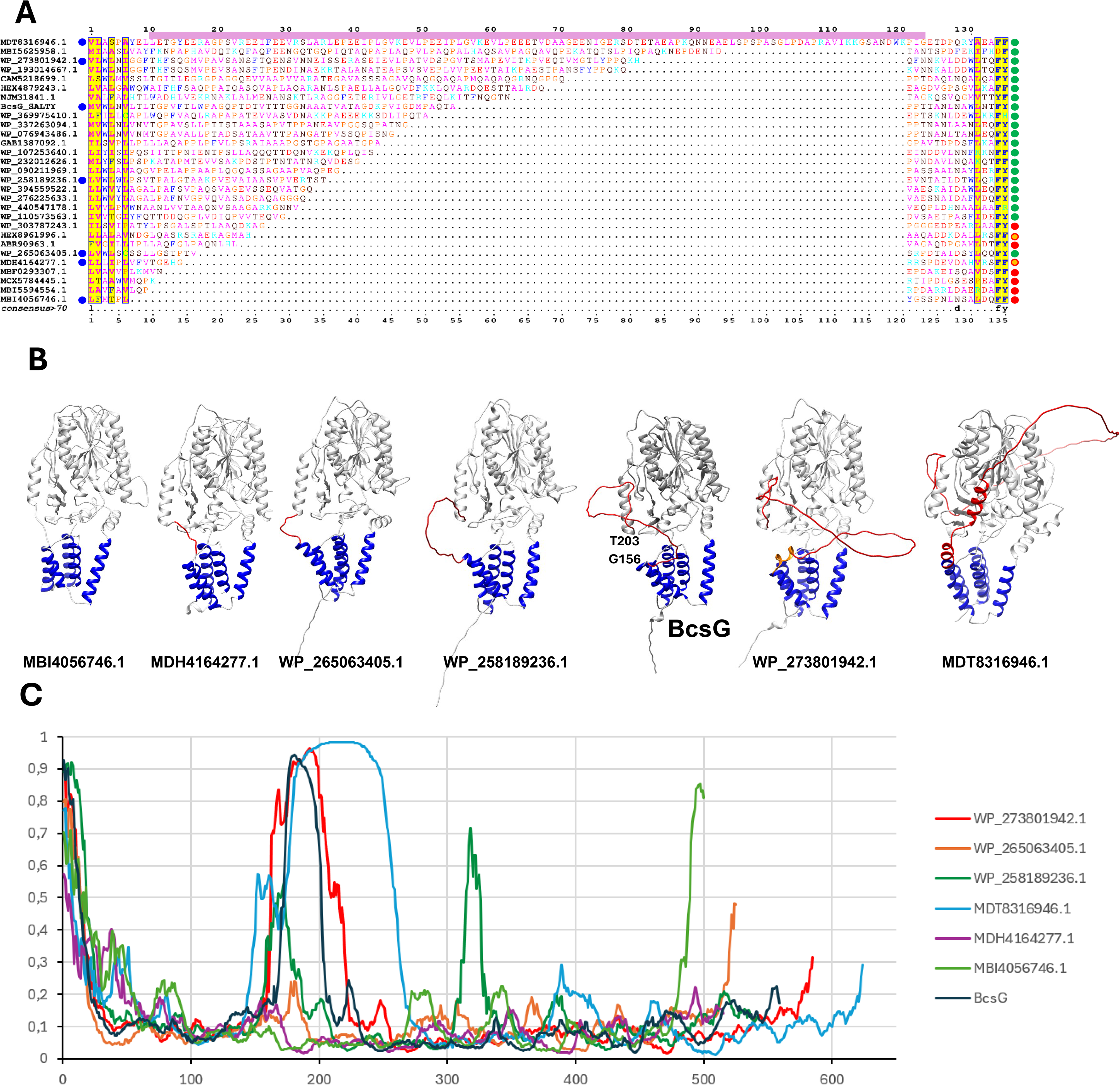
Length spectrum of N-Linker sequences of select BcsG homologs and their AlphaFold 3 models. (**A**) N-Linker sequences of selected BcsG homologs from longest to shortest identified linker sequence. Species assignment is indicated in Table S1. Indicated by a blue dot (left of alignment) are BcsG homologs displayed as structural models in (**B**). Association with a *bcs* gene cluster (as indicated by the co-localisation of *bcs* genes +/- 5 genes in the respective contig) is indicated by a green dot (right of alignment), no indication of *bcs* gene cluster colocalization with *bcsG* is indicated by a red dot and uncertain *bcsG* localization in indicated by an orange dot. (**B**) AlphaFold 3 models of selected BcsG homologs displaying different N-linker lengths. In dark blue, transmembrane helix; in red, N-linker connecting the 4^th^ or 5^th^ transmembrane helix with the α-helix of the linker. (**C**) Prediction of unstructured regions of BcsG and homologs from (**B**) by AIUPred.

### BcsG encoding genes with very short linkers are not co-localized with cellulose biosynthesis genes

Construction of a BcsG phylogenetic tree with the Maximum-Likelihood approach indicated the BcsG homologs can be subdivided into at least seven distinct clades, however linker length and consequently association with a *bcs* gene cluster is not correlated with distinct clades (Fig. 3; Fig. S3). Several remarkable features among BcsG homologs with respect to linker length were, however, observed. While BcsG with the longest recognized linker belonged to an unknown bacterium, a BcsG homolog with a long linker extending far beyond the *S. typhimurium* linker is encoded by *Proteus* spp. genomes. Species of the genus *Proteus* including the well-investigated *Proteus vulgaris* and *Proteus mirabilis* were observed to code for two cellulose biosynthesis operons, but cellulose production has not been observed (20). The BcsG homolog is part of one of the two cellulose biosynthesis gene clusters.

**Fig 3.**
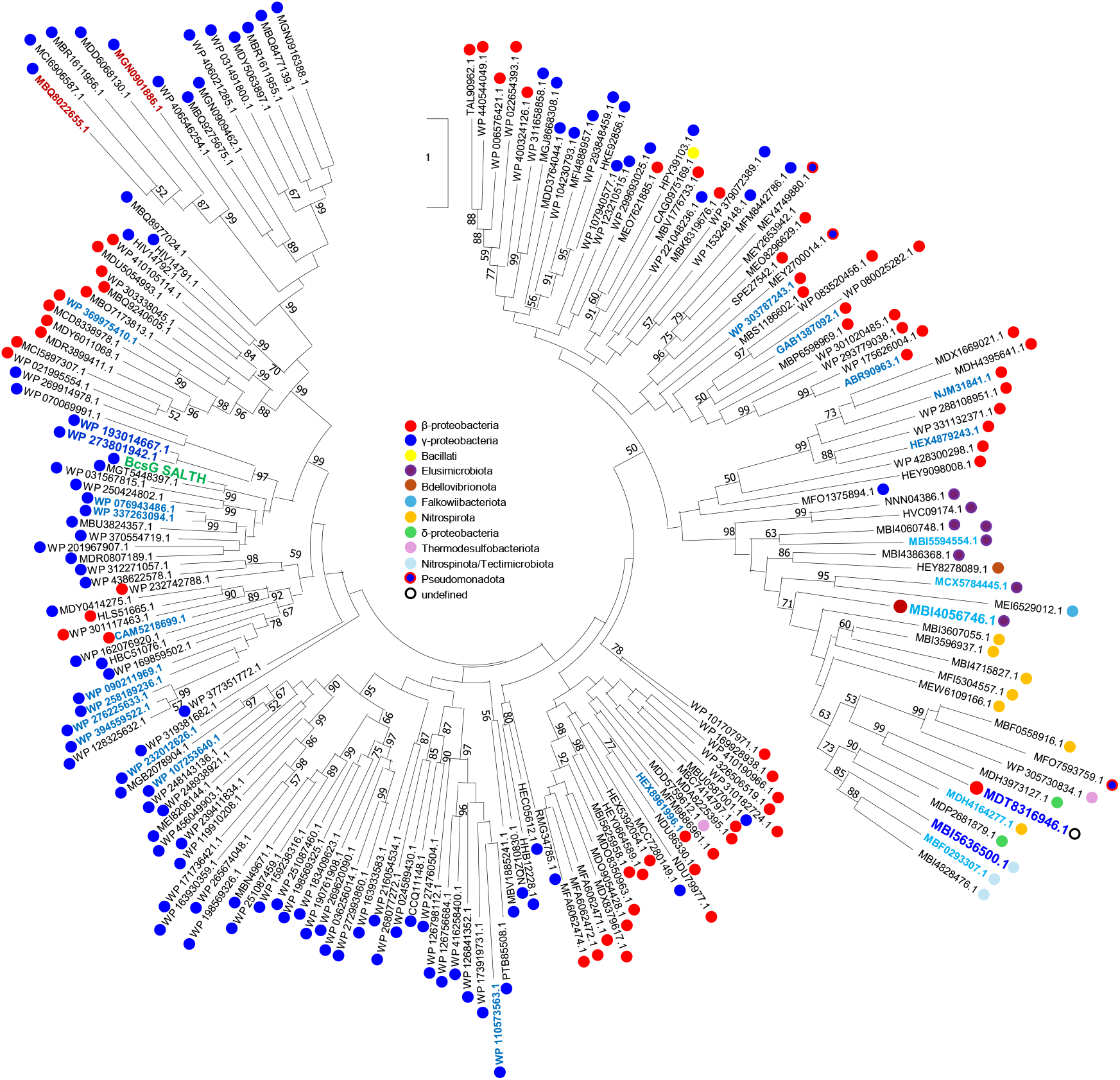
Phylogenetic tree of BcsG homologs. BcsG homologs were retrieved from the NCBI database using BcsG of *S. typhimurium* ATCC 14028 as a query for Blast search (42). Sequences were aligned with ClustalX 2.1 (43) and manually curated in GeneDoc. Sequences with >80% homology have been removed with Jalview (44) with reference sequences manually maintained. A phylogenetic tree was constructed in MEGA 11.0 (51) with Maximum Likelihood approach with1000 bootstrap iterations and the resulting tree displayed. Phylum of the BcsG homologs is indicated. Proteins with largest N-Linker length are indicated in dark blue (protein with largest linker indicated with red dot) and proteins with shortest N-linkers indicated in light blue (protein with shortest linker indicated with red dot). Proteins with linker sequences displayed in Figure 2A are indicated with intermediate blue.

Of note, BcsG homologs with the shortest N-linkers are not directly associated with a cellulose biosynthesis gene cluster (Fig. 2A-C; Fig. S2A; Table S1). species with ‘free-standing’ BcsG with short N-linkers belong to the *Elusimicrobiota*, *Jantinobacterium* and *Nitrospirinae* genera. We cannot say with 100% certainty that cellulose biosynthesis gene clusters are not encoded elsewhere on the genome as the genome sequences are not complete. Furthermore, we identified BcsG homologs that clearly lack the catalytic amino acids indicating lack of transferase activity. Also, although Gram-positive bacteria can synthesize cellulose, BcsG seems to be restricted to Gram-negative bacteria.

### Linkers of other alkaline phosphatase superfamily members are distinct and less variable

As the length of the amino acid sequence of the BcsG N-linker is highly variable, we were wondering whether amino acid sequence and length variability is a specific features of BcsG or whether other alkaline phosphatase superfamily members display similar linker variability. To this end, investigating homologs of the other six alkaline phosphatase superfamily members of *S. typhimurium* ATCC 14028, EptA, EptB, CptA, OpgB, OpgE, PbgA and LtaS of *Staphylococcus aureus* as a well-investigated Alkaline Phosphatase superfamily member as representative of a Gram-positive species as a query, we investigated the N-linkers of up to 5000 representative non-redundant homologs starting with each of the proteins as a query. Remarkably, N-linker length was significantly more conserved in the alkaline phosphatase superfamily members even down to homologs with a similarity of 30% or less (Fig. S4). As exemplified with OpgE, PbgA and LtaS, which transfers phosphoglycerol phospholipid headgroups to the lipoteichoic acid (LTA), a component of the cell wall of Gram-positive bacteria the linker sequences of even distantly similar homologs are significantly more congruent (Fig. S4). Thus the high flexibility in length and the unstructured ‘random coil’ structure of the 5’ part of the linker of BcsG, the N-linker, is unique among the alkaline phosphatase superfamily members of *S. typhimurium* and beyond.

### The N-linker length is negatively correlated with transfer activity and/or cellulose biosynthesis

As the linker length was observed to be substantially variable among BcsG homologs, we were wondering about the physiological impact of different linker length and the functional flexibility of the BcsG N-linker with the *S. typhimurium* protein as an example. To this end, we constructed a number of N-linker deletions starting with deletion of the first 18 N-terminal and the last 17 C-terminal amino acids while leaving the first ten and last two amino acids of the N-linker sequence, GPVFTLWPAG and PT, respectively untouched as a frame to ensure domain flexibility (Fig. 2 and 4A; Fig. S2A). In addition, these first ten amino acids were identified to form an α-helix in some of the BcsG homologs indicating that the fifth transmembrane helix might be part of a locally flexible α-helix even as the amino acids at the N-terminal start of the N-linker, glycine and proline, are helix breaking.

**Fig 4.**
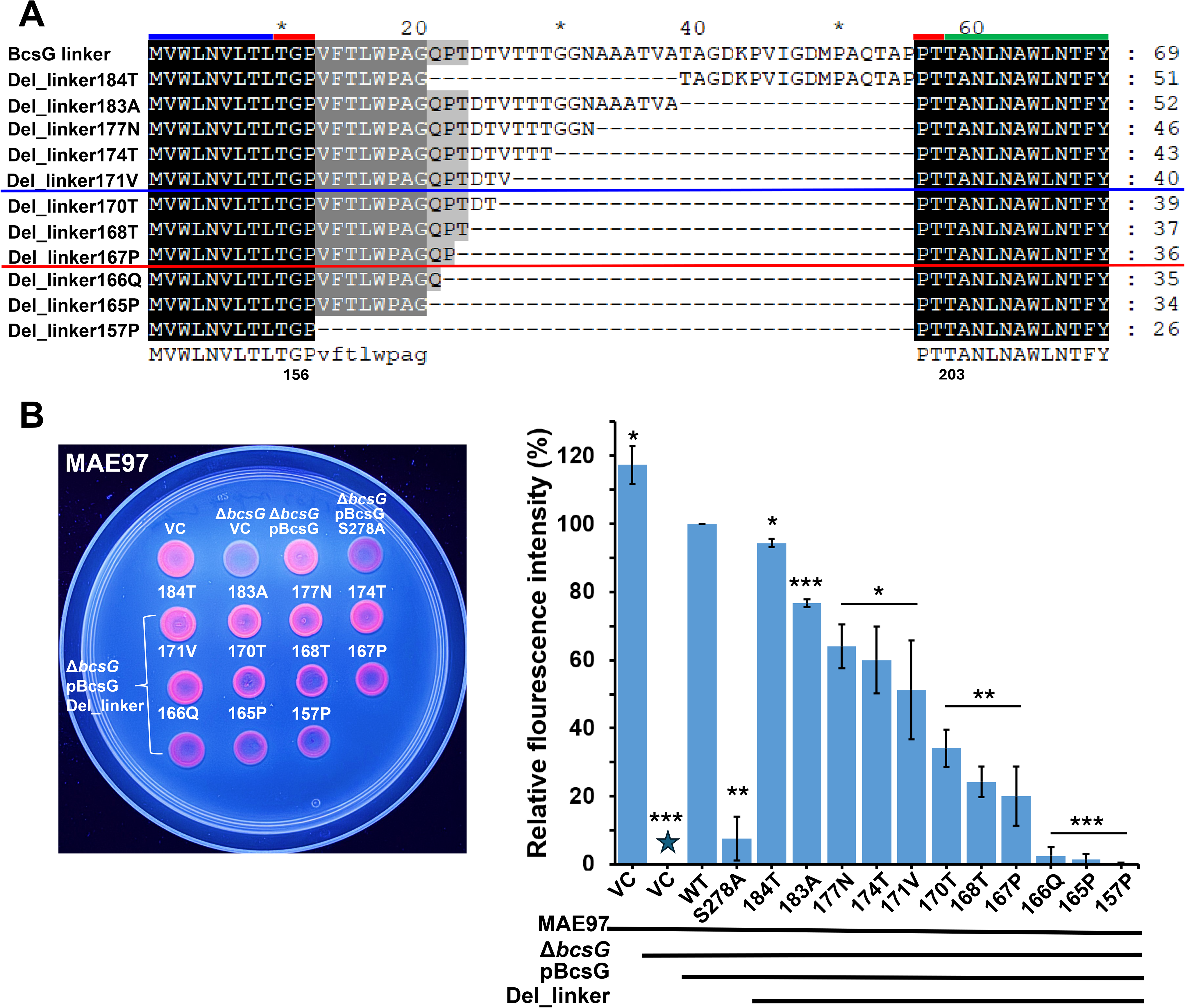
Effect of N-linker length of BcsG on pEtN transfer to the glucan chain synthesized by the cellulose synthase BcsA. (**A**) Display of N-linker variant constructs for BcsG. Underlaid in black are invariable sequences. Red bars above sequences indicate maintained N-linker sequences; blue bar indicates sequence of fifths transmembrane α-helix; green bar indicates linker α-helix. Above blue line indicates N-linker BcsG variants with >50% CR-based fluorescence output compared to wild type BcsG. Above red line indicates <20% CR-based fluorescence output compared to wild type BcsG. (**B**) Congo red fluorescence of *S. typhimurium* MAE97 Δ*bcsG* colonies complemented with wild type BcsG and its different N-linker variants (left). MAE97 vector control (VC) and MAE97 Δ*bcsG* pBcsG were positive, while MAE97 Δ*bcsG* VC and MAE97 Δ*bcsG* pBcsG S278A complemented with catalytically inactive BcsG were negative control references. VC=pBAD30. pBcsG=BcsG cloned in pBAD30 under the control of the AraC repressor and the L-arabinose promoter (2). Right; Quantification of CR fluorescence after incubation on LB without NaCl agar plates at 37□ for 8 h. Data are based on three biologically independent experiments.

Functionality of the enzymatic transfer of the phospholipid headgroup pEtN to the glucan chain was assessed by selective Congo red fluorescence after excitation of CR agar plate growth colonies with UV light of 365 nm (21). Congo red fluorescence indicates the degree of pEtN substitution of the glucan chain. Deletion of the first 18 N-terminal and the last 17 C-terminal amino acids of the N-linker sequence reduced apparent pEtN transfer to the glucan chain by 3 and 25%, respectively, compared to the full length protein although 38 and 35% of the N-linker sequence has been removed (Fig. 4B). With the first 18 N-terminal amino acids of the linker in place, gradually deleting additional amino acid from the C-terminal end retained >50% CR fluorescence upon deletion of in total 29 C-terminal amino acids. While deletion of four additional amino acid retained >20% CR fluorescence, deletion of additional amino acids including VFTLWPAG leaving only two N-linker amino acids substantially reduced pEtN-modified glucan chain based CR fluorescence (Fig. 4B). Concomitantly with the Congo red fluorescence, the roughness of the colony on the Congo red agar plate gradually diminished (Fig. S5A). Assessment of Calcofluor white binding of colonies showed a gradual diminished Calcofluor white binding upon reduction of N-linker length (Fig. S5B). Colony morphology on LB without NaCl agar plates did not indicate a correlation of colony morphotype with N-linker length. (Fig. S5C).

Correlating these results with the N-linker length and chromosomal position of BcsG homologs indicated that BcsG homologs with short linkers that are associated with *bcs* gene clusters should be capable to transfer pEtN to an emerging glucan chain although potentially with variable and restricted efficiency and/or promote less cellulose biosynthesis (Fig. S5D). Remarkably BcsG homologs with a N-linker length that does not permit pEtN transfer are not associated with *bcs* gene clusters and are therefore hypothesized to possess a different substrate(s).

### N-linker length of BcsG is negatively correlated with steady state BcsG levels, but does not affect levels of the BcsA cellulose synthase

As reduced pEtN transfer efficiency was observed upon gradual shortening of the N-linker, we were wondering which mechanism might restrict transfer. Therefore, we investigated first whether the N-linker length had an effect on the steady state levels of BcsG (Fig. 5A and B). To this end, we selected two protein variants with a significantly shorter N-linker, BcsG_Δ170T_, which retained below 40% fluorescence and BcsG_Δ157_ which contains only the four amino acid long N-linker to assess protein levels compared to the full length protein. Surprisingly, the BcsG variants with the shorter linkers showed reduced protein steady state levels of 66.8 and 37.8 %, respectively compared to the full length protein.

**Fig 5.**
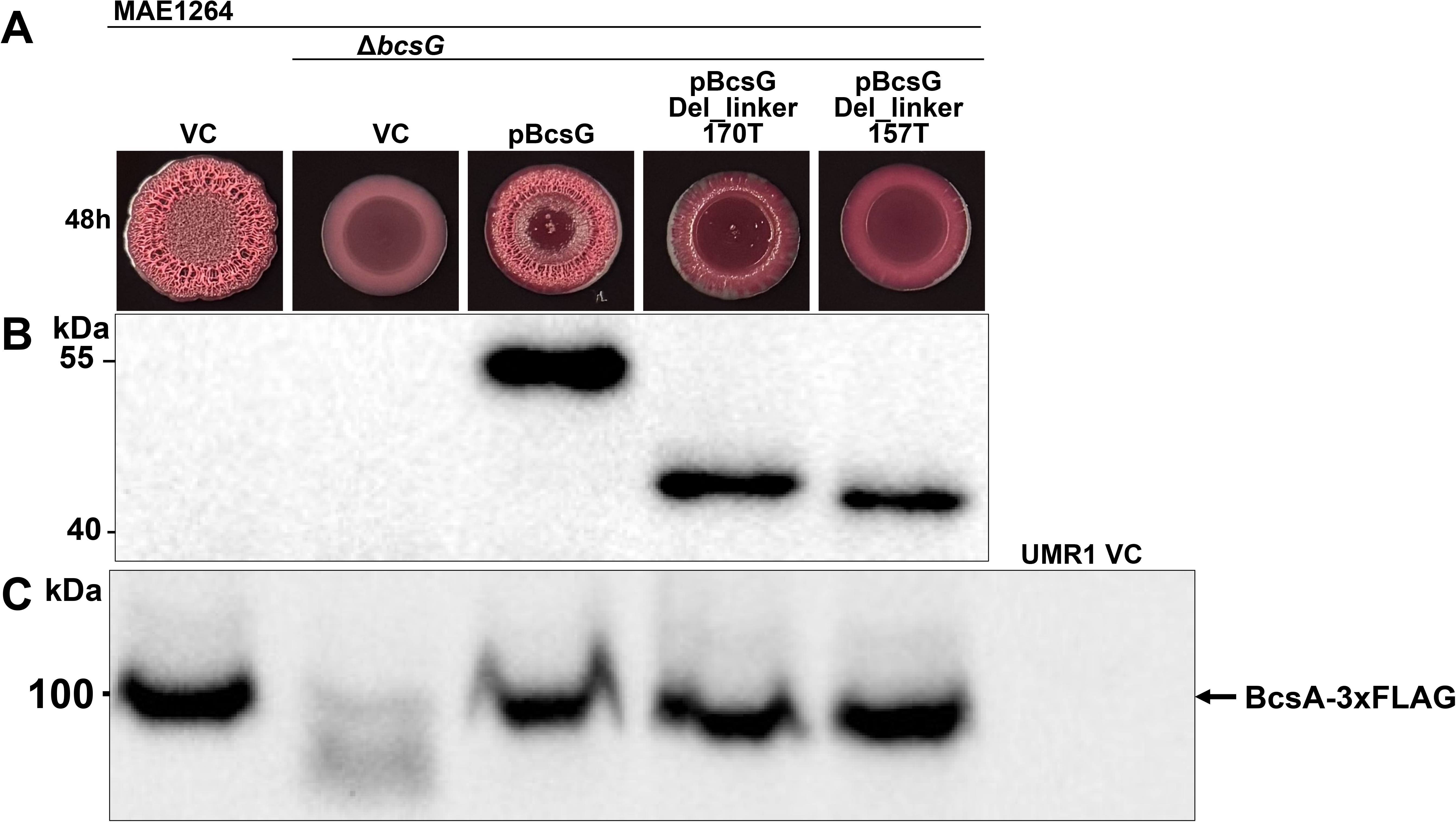
Effect of linker length on BcsG and cellulose synthase BcsA steady state levels. (**A**) Colony morphotypes of MAE1264 Δ*bcsG* complemented with selected N-linker variants BcsG Del_linker 170T and BcsG Del_linker 157T. (**B**) Steady state levels of BcsG and its variants as indicated in (**A**). Protein levels were detected by the 6xHis-tag antibody (Qiagen). (**C**) Steady state levels of BcsA in MAE1264 Δ*bcsG* complemented with selected N-linker variants BcsG Del_linker 170T and BcsG Del_linker 157T. Protein levels were detected by antibody against 3x FLAG-tag. **A**-**C**. References for comparison are MAE1264 VC and MAE1264 Δ*bcsG* pBcsG as positive controls and MAE1264 Δ*bcsG* VC as negative control. MAE1264=MAE97 BcsA-3xFLAG; VC=pBAD30. Bacterial cells were incubated on LB without NaCl agar plates at 37□ for 48 h.

We were further wondering whether protein levels of the cellulose synthase BcsA were affected by the BcsG protein variants with different N-linker length (Fig. 5C). Recently, we demonstrated that the membrane part including the linker of BcsG up to amino acid 210 is sufficient to substantially stabilize the cellulose synthase BcsA by preventing degradation and/or aiding membrane insertion (2). In accordance with our previous findings, BcsA was not affected in its steady state levels in the respective BcsG short N-linker mutants despite of lower BcsG levels. This finding might indicate that BcsG concentrations upon overexpression are significantly higher than BcsA concentrations in the membrane upon expression from the chromosomal promoter leaving the significant reduction of BcsG insignificant to prevent BcsA-BcsG interaction (22).

### The phosphatidylethanolamine content of the membrane can affect BcsG transfer activity

Although the phosphorylethanolamine decorated phospholipid is the most frequent phospholipid in the inner membrane, the phosphorylethanolamine decorated phospholipid is unequally distributed between the outer and inner leaflet of the inner membrane constituting only approx. 25% of the phospholipids in the outer leaflet (23). It is hypothesized that the phosphorylethanolamine phospholipid headgroup is collected by the periplasmic catalytic domain from the molecules of the inner leaflet and transferred to the growing glucan chain. We were therefore wondering whether the phosphatidylethanolamine content restricts the efficiency of the pEtN transfer. Phosphatidylethanolamine is synthesized from the phosphatidylserine phospholipid intermediate (Fig. 6A). PS phospholipid is synthesized from CDP-1,2-diacyl-sn-glycerol and L-serine by the PssA CDP-diacylglycerol-serine O-phosphatidyltransferase (Fig. 6A). As the phosphatidylserine phospholipid concentration in the membrane is low, overexpression of PssA should increase phosphatidylethanolamine levels. On the other hand, phosphatidylglycerol is the second most common phospholipid in the inner membrane. PG phospholipid is synthesized from CDP-1,2-diacyl-sn-glycerol and sn-glycerol 3-phosphate upon catalysis by PgsA CDP-diacylglycerol--glycerol-3-phosphate **3-**phosphatidyltransferase. As the same substrate CDP-1,2-diacyl-sn-glycerol is used, PG phospholipid synthesis is directly competing with the synthesis of the phosphatidylethanolamine.

**Fig 6.**
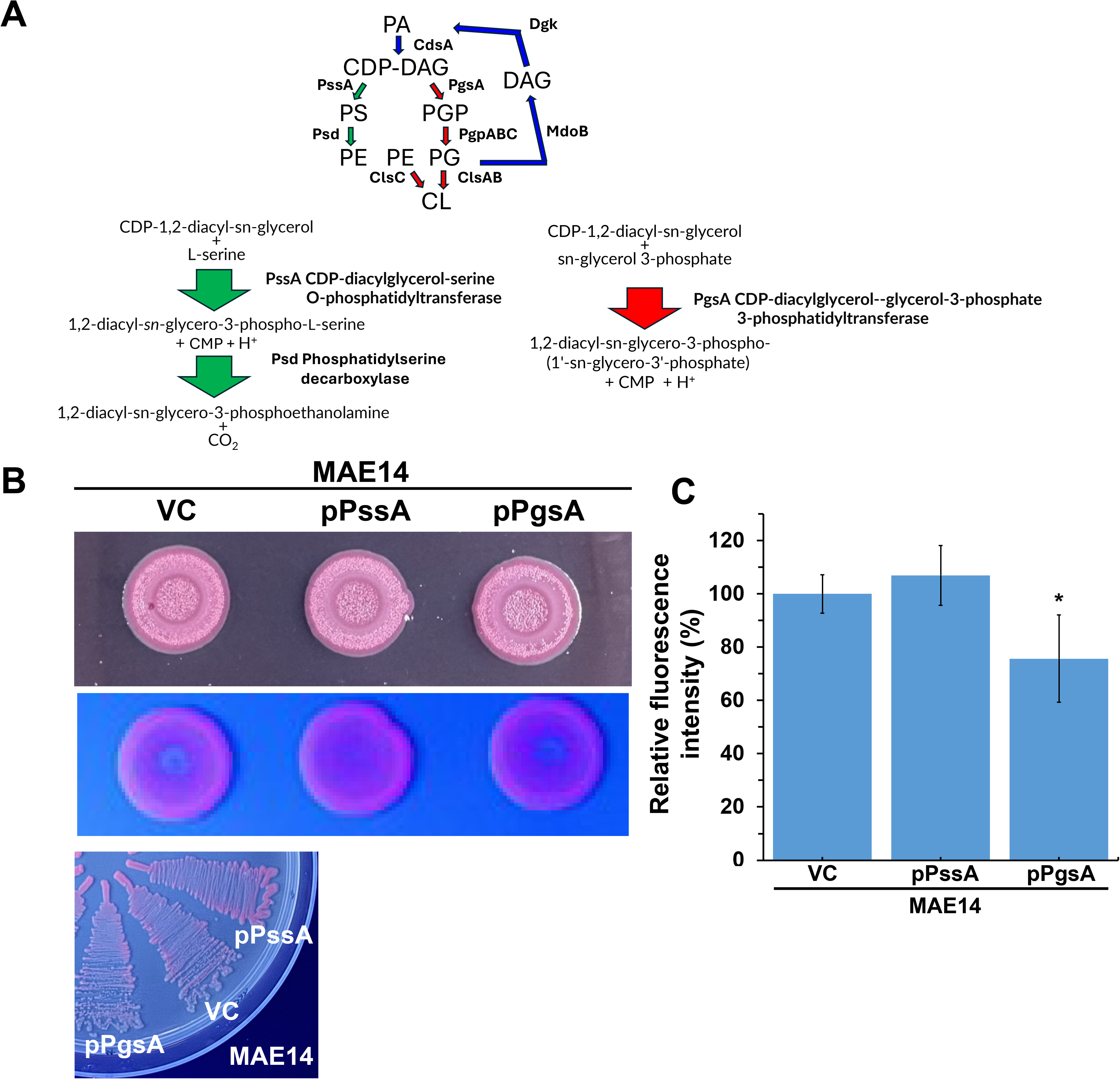
Effect of overexpression of PssA and PgsA on pEtN transfer efficiency to the glucan chain synthesized by the cellulose synthase BcsA in *S. typhimurium* MAE14 displaying regulated cellulose biosynthesis. (**A**) Pathway leading to the biosynthesis of phosphatidylethanolamine and phosphatidylglycerol. Abbreviation for molecules: PA=phosphatidate; PS=phosphatidylserine; PG=phosphatidylglycerol; PE=phosphatidyl-ethanolamine; CDP-DAG=CDP-1,2-diacyl-sn-glycerol; DAG=1,2-diacyl-sn-glycerol; CL=cardiolipin. Enzymes involved: CdsA=phosphatidate cytidylyltransferase; PssA=CDP-diacylglycerol-serine O-phosphatidyltransferase; Psd=phosphatidyl-serine decarboxy-lase; PgsA=CDP-diacylglycerol-glycerol-3-phosphate 3-phosphatidyl-transferase; PgpABC=phosphatidylglycerophosphatase A, B and C; ClsAB=cardiolipin synthase A and B; ClsC=cardiolipin synthase C; Dgk=diacylglycerol kinase; MdoB (OpgB)=-phosphoglycerol transferase I. (**B**) Effect of overexpression of PssA and PgsA on the colony morphology and CR fluorescence of *S. typhimurium* MAE14 on CR and CRonly agar plates. Colonies were grown for 72 h on CR and CRonly agar plates to assess colony morphology and CR fluorescence, respectively. Streaks were grown for 24 h on CRonly agar plates to assess fluorescence. MAE14=UMR1 Δ*csgBA*; VC=vector control pBAD30; pPssA and pPgsA are cloned in pBAD30. (**C**) Quantification of the effect of overexpression of PssA and PgsA on the Congo red fluorescence of MAE14. Data are presented as mean ± SD from three independent biological replicates (n = 3). *=p<0.05 by student’s t-test compared to MAE14 VC.

We were therefore overexpressing PssA and PgsA to observe their effects on phosphorylethanolamine transfer to the glucan chain. In order to sensitively observe differences in pEtN transfer we used MAE14, a derivative of the wild type *S. typhimurium* UMR1, as the host strain which displays highly regulated cellulose expression. To this end, we observed that expression of PssA showed enhanced fluorescence, while overexpression of PgsA displayed reduced fluorescence indicating reduced pEtN transfer due to reduced availability of phosphatidylethanolamine (Fig. 6B and C). These observations are consistent with the hypothesis that the concentration of phosphatidylethanolamine is enhanced by overexpression of PssA, while overexpression of PgsA reduces the phosphatidylethanolamine content.

### An N-linker longer than in BcsG from *S. typhimurium* has no apparent effect on transfer activity

As insufficient phosphatidylethanolamine availability might be one restriction that limits pEtN transfer, we aimed to assess the role of a linker longer than in BcsG_St_. To this end we choose the BcsG_Pv_ linker from *P. vulgaris* which is with 78 amino acids 23 amino acids longer than the BcsG_St_ linker (Fig. 7; Fig. S6). As the membrane part of BcsG_St_ was shown to interact with and stabilize the cellulose synthase BcsA, a hybrid protein was assembled with BcsG_St_ as the scaffold. In the hybrid proteins the linker of BcsG_St_ from amino acid G156 to T202 was replaced with the linker of BcsG_Pv_ from amino acid G149 to N226 (G_149_GFTHFSQGMVPAVSANSFTQENSVNNEISSERASEIEVLPATVDSPGVTSMAPEVITKPVEQTVMGTLYPPQKHQFN_226_; (Fig. 7)). We observed that this hybrid protein transferred phosphorylethanolamine with a similar efficiency as the wild type BcsG as judged by the Congo red fluorescence assay.

**Fig 7.**
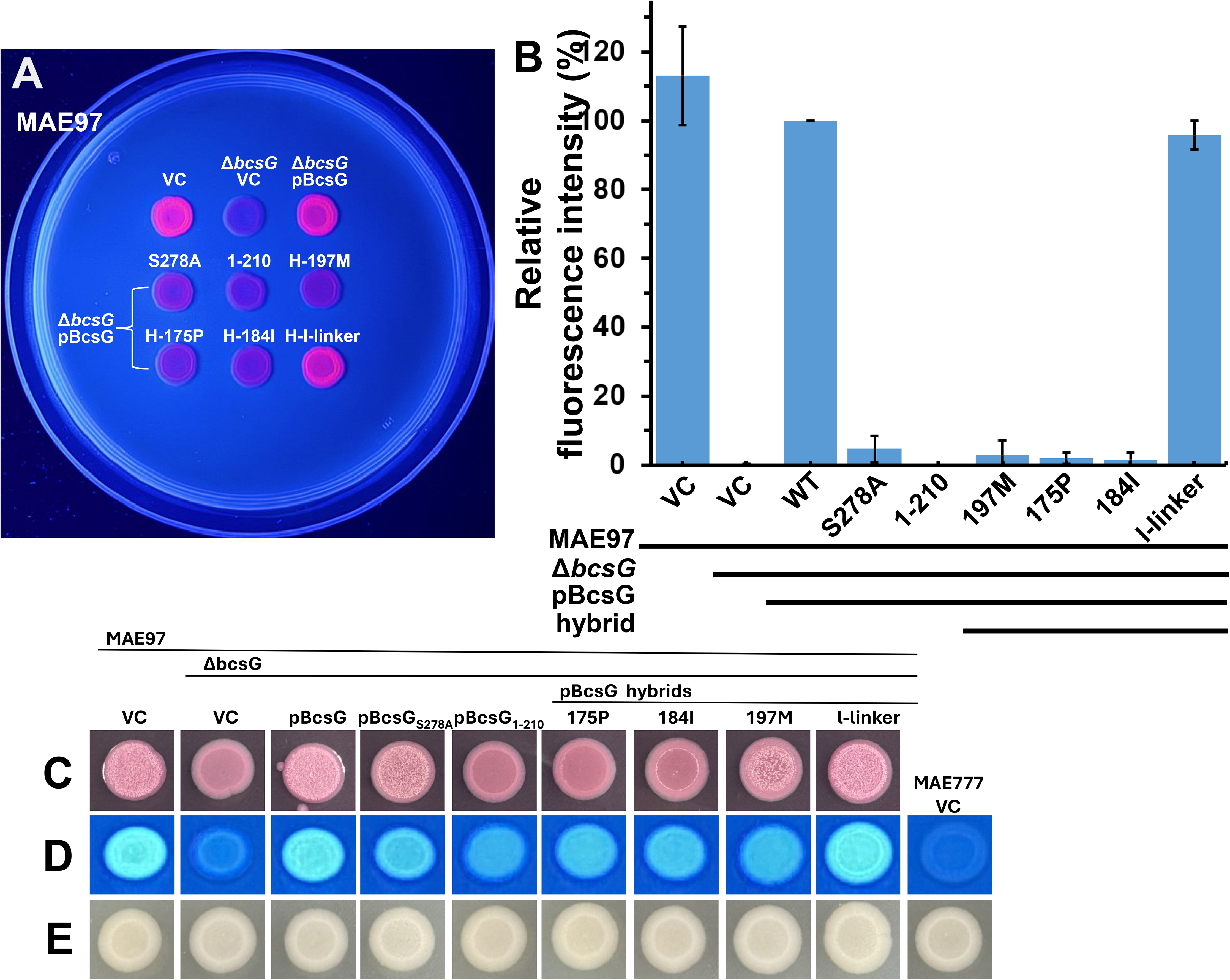
Effect of N-linker and catalytic domain exchange on the pEtN transferase activity of BcsG and OpgE. (**A**) Congo red fluorescence of *S. typhimurium* MAE97 Δ*bcsG* colonies complemented with wild type BcsG, the BcsG-linker hybrid and the BcsG-OpgE hybrid proteins. (**B**) Quantification of CR fluorescence. Data are based on three biologically independent experiments. (**C**) Colony morphology of *S. typhimurium* MAE97 Δ*bcsG* colonies complemented with wild type BcsG, the BcsG-linker hybrid and the BcsG-OpgE hybrid proteins on Congo red agar plates. Cells were grown on LB without NaCl agar plates at 37□ for 24 h. MAE97 vector control (VC) and MAE97 Δ*bcsG* pBcsG were positive, while MAE97 Δ*bcsG* VC, MAE97 Δ*bcsG* pBcsG_1-210_ and MAE97 Δ*bcsG* pBcsG S278A complemented with catalytically inactive BcsG were negative control references. VC=pBAD30. pBcsG=BcsG cloned in pBAD30 under the control of the AraC repressor and the L-arabinose promoter (2).

### The catalytic domain of OpgE has an inconclusive effect on modification of the 1,4-β-D glucan chain

Not only BcsG, but also EptA, EptB, CptA and OpgE conduct transfer of pEtN, however to molecules of different structure or linkage such as the core part of LPS and to osmoregulated periplasmic glucans. Alkaline phosphatase superfamiliy members can, however, be promiscuous with respect to the substrate. For example, Cj0256 of *Campylobacter jejuni* has been shown to transfer pEtN not only to the core of the lipooligosaccharide, but also to the flagellar rod protein FlgG (4). We were therefore wondering whether spatial distance is (partly) determining the substrate specificity of Alkaline Phosphatase superfamily members. In *S. typhimurium*, most closely related to BcsG with respect to substrate specificity and catalytic activity is OpgE which transfers pEtN to covalently link the molecule to the 2’OH group of glucose in 1,2-1,6-beta-linked osmoregulated periplasmic glucan to form a phosphodiester bond. Furthermore, OpgE has a linker structure most closely resembling BcsG_St_, although the N-linker is highly reduced (Fig. S1). The N-terminal part, the unstructured N-linker is with 12 amino acids shorter than in BcsG_St_, while the alpha-helical secondary structures are extended. To this end, three different hybrid proteins with the membrane part and the N-linker of BcsG_St_ were assembled. In a first construct the N-terminal membrane part and the entire N-linker until T202 were fused to the C-terminal part of OpgE starting from P175, I184 and M197 (Fig. 7; Fig. S6). Complementation of the S. typhimurium MAE97 Δ*bcsG* with the three hybrid proteins did, however, not trigger pEtN transfer as assessed by the Congo red fluorescence assay (Fig. 7C). We observed, however, that the colony morphotype on CR agar plates did not maintain the smooth pink morphotype as upon production of unmodified cellulose, but gradually displayed rough colony morphology. This wrinkled morphoptype was most pronounced upon expression of the BcsG-OpgE_T202-M197_ hybrid protein, while it was only marginally observed with high spatial restriction as a defined ring upon expression of BcsG-OpgE_T202-I184_ hybrid protein and barely visible with BcsG-OpgE_T202-P175_. Thus we hypothesize that the BcsG-OpgE_T202-M197_ hybrid protein might efficiently modify the glucan chain, however, not at the C6-OH position. Such a modification might therefore not cause a distinct Congo red fluorescence.

## DISCUSSION

In this work we show that the length of the unstructured N-linker that connects the fifth transmembrane helix with the catalytic alkaline phosphatase superfamily domain of BcsG has a functionality, unsurprisingly, in the transfer efficiency of the phosphorylethanolamine (pEtN) headgroup and potentially in the amount of synthesized cellulose, and, surprisingly, is positively correlated with protein stabilization. Furthermore, our results indicate that spatial proximity to the cellulose synthase BcsA can (partially) determine substrate specificity.

Our analysis of the role of the BcsG linker was initiated as the role and functionality of (unstructured) linker sequences is poorly defined. Linker sequences can be entities that allosterically transfer signals between different domains. Such linkers conventionally possess an α-helical secondary structure and can dimerize upon signal perception to form a conditional coiled-coil structure (24, 25). Other signal transduction modules such as HAMP constitute a domain on its own (26). However, linkers can play a more determinative role in protein functionality beyond solely providing a ‘random coiled’ structure that provides structural flexibility and ligand binding promiscuity (27–30). For example, the amino acid composition and flexibility of the linker between the second transmembrane domain and the cytoplasmic catalytic domains of the two FtsH proteases is determinative for substrate processing in *Pseudomonas aeruginosa* clone C (31).

In the case of BcsG, the full extent of linker, and in particular N-linker functionality is currently unknown. Eventually, combining the bioinformatic and experimental data different scenarios arise with respect to pEtN transfer to the growing glucan chains synthesized by the cellulose synthase BcsA (Fig. 8). The N-terminal part of the linker of BcsG homologs which connects the fifth transmembrane helix with the periplasmic catalytic domain is highly flexible in length and diverse in amino acid sequence although amino acid such as alanine, threonine, serine, proline and glycine are predominantly present (Fig. S 1B). BcsG homologs with a very short N-linker of up to 20 amino acids are frequently not co-localized with cellulose biosynthesis operons as indicative by the presence of *bcsA* or other genes indicative for a cellulose biosynthesis gene cluster suggesting a function different from the transfer of pEtN to a 1,4-β-D-glucan chain. This finding is grossly congruent with findings in this work where BcsG_St_ variants possessing a N-linker with less than 18 amino acids inefficiently transfer pEtN to the glucan chain (Fig. 3).

**Fig 8.**
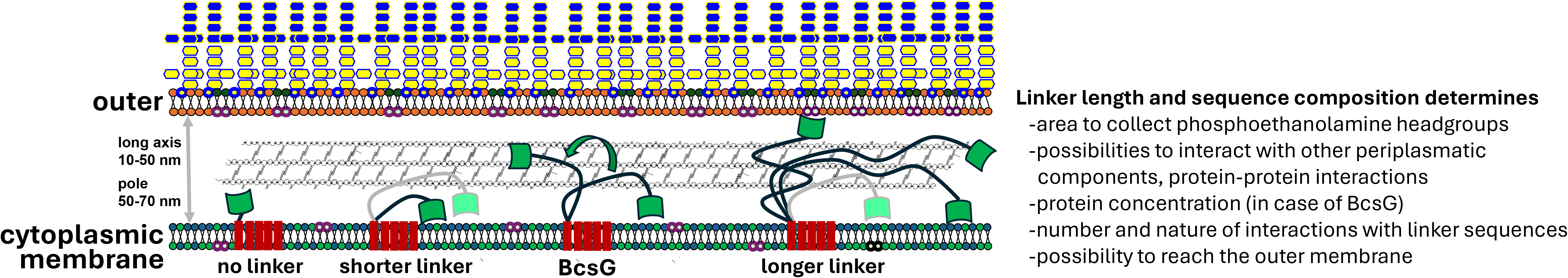
Consequences of different N-linker length of BcsG.

BcsG homologs with short N-linkers of as less than 20 amino acids can be associated with a cellulose biosynthesis operon and are hypothesized to transfer a phosphorylethanolamine headgroup albeit with lower efficiency leading to a less decorated glucan chain and/or stimulating less cellulose biosynthesis. Compensatory mechanisms that overcome a shorter linker length, however, might be in place. As a possibility, synthesis of the glucan chain by BcsA is slower in this constellation which would subsequently enable an equally densely decorated glucan chain. Also, the concentration of phosphoryl-pEtN in the membranes might be higher which might enable a more efficient collection of pEtN phospholipid headgroups. Last, but not least modulation of the degree of pEtN modification of the glucan chain might allow the synthesis of cellulose macromolecules with distinct properties by combination of different mechanisms modulating the degree of pEtN decoration.

A shorter linker length might, however, restrict transfer of the pEtN phospholipid headgroup in more than one way. Firstly, the area of collection of pEtN phospholipid headgroups from the outer leaflet of the cytoplasmic membrane is restricted to a radius defined by the linker length with the phosphoryl-pEtN content of the outer leaflet only 25% of entire cytoplasmic membrane (23). Second, although the source of the pEtN phospholipid headgroups is currently unknown, a short linker will prevent to harvest pEtN phospholipid headgroups from the inner leaflet of the outer membrane in addition to the harvesting from the outer leaflet of the cytoplasmic membrane (which is considered the default source of pEtN). Approximately 25% of total phosphatidyl-pEtN is constituting the inner leaflet of the outer membrane (32, 33) The ability to harvest from the outer membrane might be dependent on the width of the periplasm which is estimated to conventionally have a width between 50-70 nm at the pole and a width of approx. 10-50 nm at the lateral length of the rod and the corresponding location of the cellulose biosynthesis nanomachine. The length of the 47 aa long N-linker of BcsG is at most 19 nm (calculated with a maximum contour length of 4 Å per amino acid (34)). Third, a longer linker might enable to more flexibly approach the growing glucan chain at different positions. In this context it is not clear how the peptidoglycan chains provide space for the cellulose biosynthesis machinery and the potential flexibility for the BcsG linker. Although BcsG homologs with longer linkers might reach the inner leaflet of the outer membrane to harvest pEtN, but also have a much larger membrane area of the outer leaflet of the inner membrane to harvest pEtN headgroups, we did not notice an extended efficiency of pEtN transfer of the BcsG hybrid with a longer linker (Fig. 7).

Besides the basic functionality of pEtN transfer to the glucan chain, longer linkers provide extended possibilities for BcsG to interact with other periplasmic components such as small molecules, other non-proteinaceous periplasmic components and by protein-protein interactions. This feature prepares also for extended catalytic options reaching alternative molecules or proteins that can serve as substrates/recipients of the pEtN phospholipid headgroup. Those proteins might not be directly co-localized with BcsG or freely diffusible in the periplasm. Longer linkers extend also the regulatory options such as protein stabilization (as it seems the case for BcsG_St_) or e.g. processing by proteinases (as it is the case for OpgB and LtaS (13, 35)). We have, however, not observed that the periplasmic catalytic domain of BcsG is cleaved off (Fig. 5). In conclusion, the N-linker length might not necessarily be linearly correlated with pEtN transfer efficiency as alternative functionalities might modulate the discussed direct association between linker length and pEtN decoration

Also other well-investigated alkaline phosphatase superfamily members of *S. typhimurium* possess linkers with homologs to possess linkers of variable length. In the case of Mcr1 homologs to target the linker might be a novel approach to convert the resistance against polypeptide antibiotics targeting the outer membrane (36, 37).

## MATERIALS AND METHODS

### Strains and growth conditions

The K-12 derivative *Escherichia coli* TOP10 has been the host strain for cloning. *E. coli* TOP10 (without and with inserted plasmids) was grown on LB agar plates or in LB medium at 37°C, supplemented with the required antibiotics. *Salmonella typhimurium* MAE52 (ATCC14028 P*csgD1* Nal^r^ rdar_28°C/37°C_ (38)). UMR1 (ATCC14028 Nal^r^ rdar_28°C_) and derivatives thereof were maintained on LB medium agar plates or Congo red (CR) agar plates (LB medium without NaCl, 40 µg/ml Congo red, 20 µg/ml Coomassie Brilliant Blue G-250. The antibiotic ampicillin was added at 100 µg/ml and induction of protein production was achieved with 0.1% L-arabinose. Strains are listed in Table S2

### Gene cloning

Open reading frames with the native 5’ Shine-Dalgarno sequence (adding 20 bp 5’ of the start codon) were cloned into pBAD30 by in vivo recombination cloning (39, 40). In short, the open reading frame and the vector were amplified with at least 15 bp overlapping primers mimicking insertion into the XbaI and SphI sites. The two linear fragments were transformed into a chemocompetent *E. coli* K-12 derivative. Primers are listed in Table S3 and plasmids in Table S4.

### Construction of linker deletions

BcsG cloned in pBAD30 has been used as template (2). Linker deletions were constructed by designing primers that overlap at the 5’ end by at least 15 bp and contain the complementary sequence to amplify the linear plasmid at the 3’ end. Overlapping 5’ ends were designed to result in *bcsG* linker deletions of variable lengths. The linear fragment was transformed into a chemocompetent *E. coli* K-12 derivative. Plasmid were recovered and constructs confirmed by sequencing. Primers are listed in Table S3 and plasmids in Table S4.

### Construction of hybrid genes

BcsG in pBAD30 has been used as template (2). Primers which overlap at the 5’ end by at least 15 bp with the insert and are at the 3’ end homologous to the *bcsG* sequence were used to amplify the plasmid. The insert to replace the homologous *bcsG* sequence was amplified using primers homologous at the 5’ end with the *bcsG* sequence (or to the vector in case of replacement of the entire catalytic domain) and at the 3’ end homologous to the desired insertion sequence. Templates to create insert were *opgE* of *S. typhimurium* UMR1 and codon optimized *bcsG* of *Proteus vulgaris* (Integrated DNA Technologies). The two linear fragments were transformed into chemocompetent *E. coli* K-12 derivatives for in vivo recombination cloning. Primers are listed in Table S3 and plasmids in Table S4.

### Plasmid transformation

Plasmid constructs were transformed into chemocompetent *E. coli* TOP10 and into electrocompetent *S. typhimurium* MAE52, UMR1 and derivatives by electroporation as described (38).

### Assessment of cellulose production on Congo red plates

Ten µl of a 2 OD_600_ suspension was spotted onto Congo red agar plates. Congo red agar plates were incubated at 37°C for up to 72 h to assess development of cellulose and pEtN modified cellulose. Positive controls were *S. typhimurium* MAE97 or MAE97 Δ*bcsG* pBcsG and negative control was MAE97 Δ*bcsG* pBAD30 empty vector control. Alternatively, S. typhimurium MAE14 (UMR1 Δ*bcsA*) was used as a host. Colony morphology and coloration was documented in a gel documentation chamber using a conventional mobile phone,

### Assessment of pEtN modified cellulose by Congo red fluorescence

Phosphorylethanolamine modification of cellulose was assessed by Congo red fluorescence (21). Ten µl of a 2 OD_600_ suspension was spotted onto Congo red only (CRonly) agar plates with LB without NaCl as growth medium. Congo red only agar plates containing 35 μg/mL Congo red were incubated at 28 or 37°C for up to 72 h to assess the degree of pEtN modified cellulose (21). Positive control were *S. typhimurium* MAE97, MAE97 pBAAD30 empty vector control, MAE97 Δ*bcsG* pBcsG and negative control was MAE97 Δ*bcsG* pBAD30 empty vector control, MAE97 Δ*bcsG* complemented with the BcsG catalytic mutant pBcsG_S278A_ and with pBcsG_1-210_ to ensure stability of the cellulose synthase BcsA. Colony morphology and coloration was documented in a gel documentation chamber using a conventional mobile phone after visualization under UV light of 365 nm wavelength. Quantification of the fluorescence intensity was performed using ImageJ software (41). The experiments were repeated independently three times.

### Quantification the fluorescence intensity using ImageJ software

Fluorescence images were acquired using a smartphone under 365 nm UV illumination and camera settings. Images were analyzed using ImageJ software (National Institutes of Health, Bethesda, MD, USA). RGB images were split into individual color channels, and the red channel corresponding to the pink fluorescence emission was selected for quantitative analysis. A region of interest (ROI) with identical size was applied to each sample. The mean gray value was measured after background correction using an ROI placed in a non-fluorescent area. Fluorescence intensity was expressed as background-corrected mean gray value. Relatively fluorescence intensity (%) = (Fluorescence intensity of test strains) / (Fluorescence intensity of MAE97 ΔbcsG pbcsG) X 100.

### SDS-PAGE and Western blot analysis

Strains were grown on LB without salt plates with Ampicillin (50 μg/mL) and 0.1% L-arabinose for 16h at 28 □. 200 μL of urea buffer (8 M urea, 2% SDS, 11% glycerol, 62.5 mM Tris–HCl, pH 6.8) was added to 5 mg cell pellet. Samples were mixed and stored at -20 □ until use. Equal amounts of protein (typically 4-6 μL) were loaded onto a SDS-PAGE gels (6% resolving gel, 4% stacking gel). Electrophoresis was performed in running buffer for 30 minutes and 120 V for 40 minutes until the dye front reached the bottom of the gel. The gel was stained with Coomassie Brilliant Blue R-250 and destained until clear protein bands were visible. Sample volume was adusted to equal anounts of protein if required. For western blot, separated proteins were transferred onto a polyvinylidene difluoride (PVDF) membrane (Millipore) using a wet transfer system at 100 V for 2 hours. The membrane was briefely rinsed in water and incubated in the Ponceau S staining solution (0.1%, 5% acetic acid) for 15 minutes. After documentation, the membrane was washed 3 times with 1XTBS-T buffer and incubated for 1 h in blocking buffer (3% BSA in 1XTBS-T) at room temperature or overnight at 4 □. After washing three times with TBS-T (5–10 min each), the membrane was incubated 1h at room temperature with the appropriate primary antibody (1:3000 for Penta-His, 1:5000 for BcsA-3xFLAG) diluted in blocking buffer succeeded by incubation with Goat Anti-Mouse IgG (1:5000; SinoBiological) solution in 10% milk powder in TBS for 1h at room temperature. Following four additional washes with TBS-T, the protein bands were visualized using an enhanced chemiluminescence (ECL) detection reagent and captured with a chemiluminescence imaging system (ChemiDoc^TM^Touch Imaging System, BIO-RAD).

### Bioinformatic approaches

BcsG, other alkaline phosphatase superfamily members encoded by *S. typhimurium* ATCC 14028, Mcr and LtaS from *S. aureus* were used as a query to retrieve homologous proteins from the non-redundant protein, alternatively the ClusteredNR database, by BLAST search with standard parameters from NCBI (42). Protein sequences were aligned with ClustalX 2.1 (43) and manually curated in Genedoc. For some analyses, proteins with >90 or >80% identity were removed with Jalview (44). Phylogenetic trees were constructed with MEGA 7.0 (45). Designation and origin of proteins are displayed in Table S5. Alignments were displayed with ESPript 3.0 (46). Amino acid frequency was assessed in Expasy with ProtParam. Conservation of amino acids in the alignments was displayed with Weblogo 3 (47). Structural models were retrieved with AlphaFold 3 (48). Visualization, manipulation and overlay of structural models was performed in Chimera 1.8 (49). Unstructured regions were assessed with AIUPred (50).

## ACKNOWLEDGEMENTS

This work received support from the Swedish Research Council (SRC), Röntgen-Ångström-Cluster (diary number 2023-06374) and the SRC for Natural and Engineering Sciences (diary number 2022-04865). The funders had no role in study design, data collection and interpretation, or the decision to submit the work for publication.

