## Supplemental Figures 1-8; Supplemental Tables 2-4 for "Role of the unstructured N-linker of the alkaline phosphatase superfamily member BcsG in the gastrointestinal pathogen *Salmonella typhimurium*"

*Running title: BcsG N-linker in Salmonella typhimurium*

Li Li and Ute Römling*

Department of Microbiology, Tumor and Cell Biology, Biomedicum, Karolinska Institutet, SE-171 77 Stockholm, Sweden

Content

Supplementary Figure S1

Supplementary Figure S2

Supplementary Figure S3

Supplementary Figure S4

Supplementary Figure S5

Supplementary Figure S6

Supplementary Material

Supplementary Table S1. Colocalisation of selected BcsG encoding genes with core *bcs* genes

Supplementary Table S2. Strains used in this study

Supplementary Table S3. Primers used in this study

Supplementary Table S4. Plasmid used in this study

Supplementary Table S5. Proteins in phylogenetic tree Figure 3

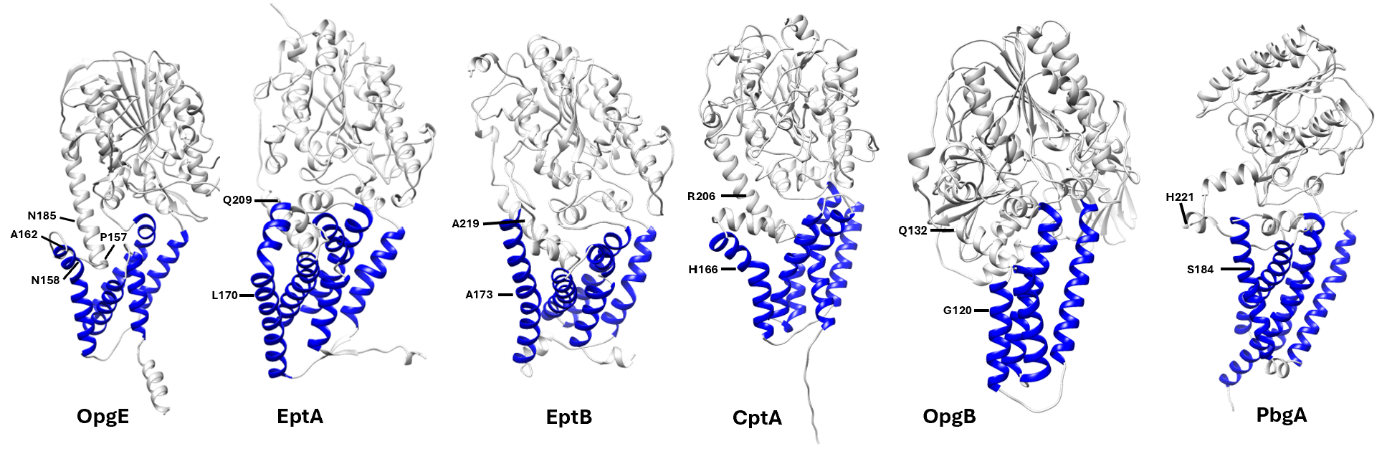

**Fig S1** AlphaFold 3 models of six of seven Alkaline Phosphatase superfamily members of *S. typhimurium* ATCC 14028. The sequence range homologous to the N-linker of BcsG (Figure 1) are indicated with the respective flanking amino acids. At the OpgE structure, an alternative sequence range that defines the N-linker sequence is indicated. In dark blue, transmembrane helix.

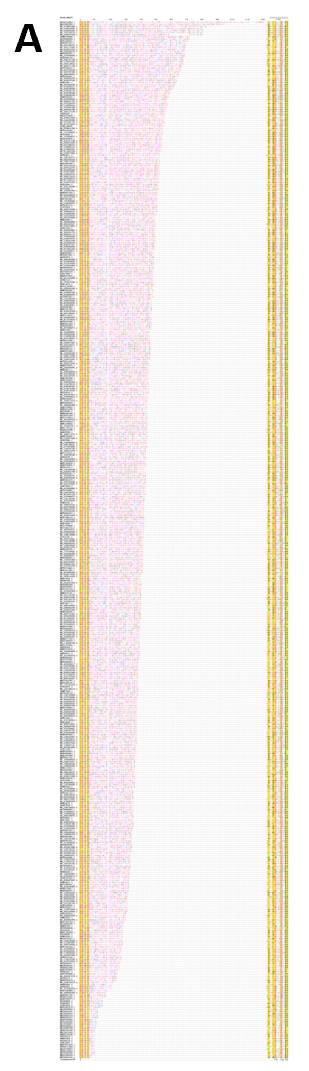

**Fig S2** Length and amino acid composition of N-linkers of BcsG homologs. (**A**) N-linkers of BcsG homologs with less than 90% sequence identity ordered according to length. Borders of linker sequences are indicated.

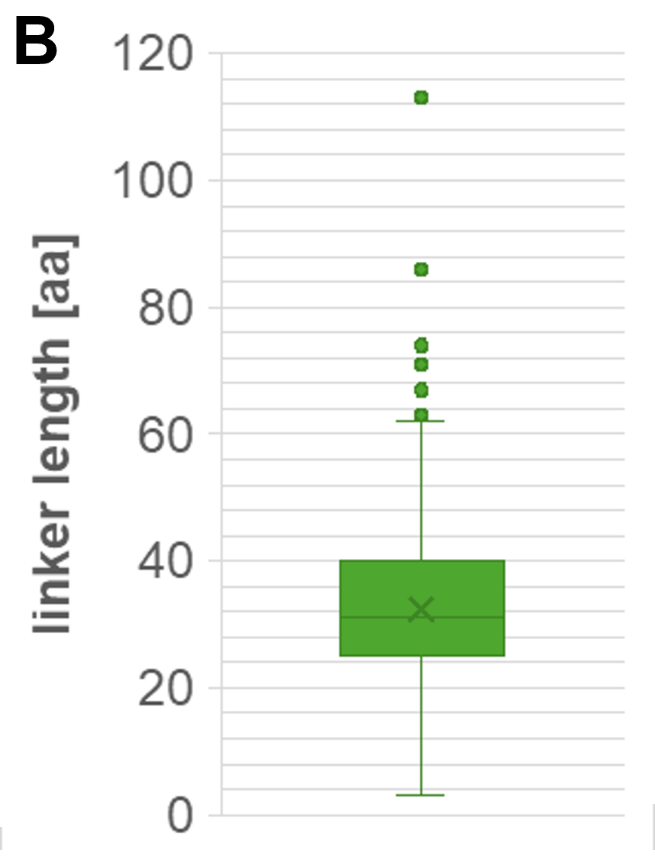

(**B**) Size distribution of linker length of BcsG homologs.

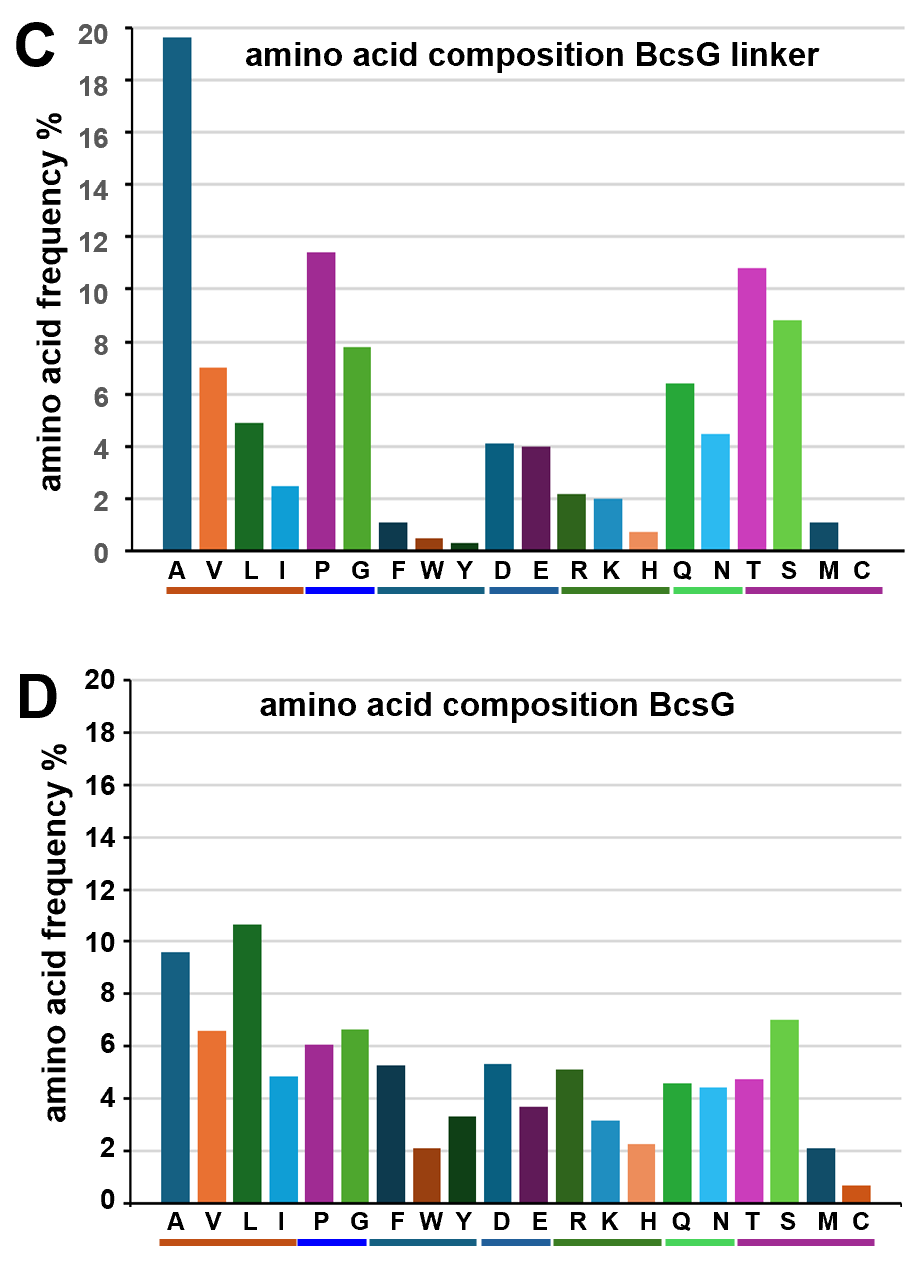

(**C**) Amino acid composition of the N-linkers based on in total 33689 amino acids. (**D**) Amino acid composition of full-length BcsG based on in total 532029 amino acids.

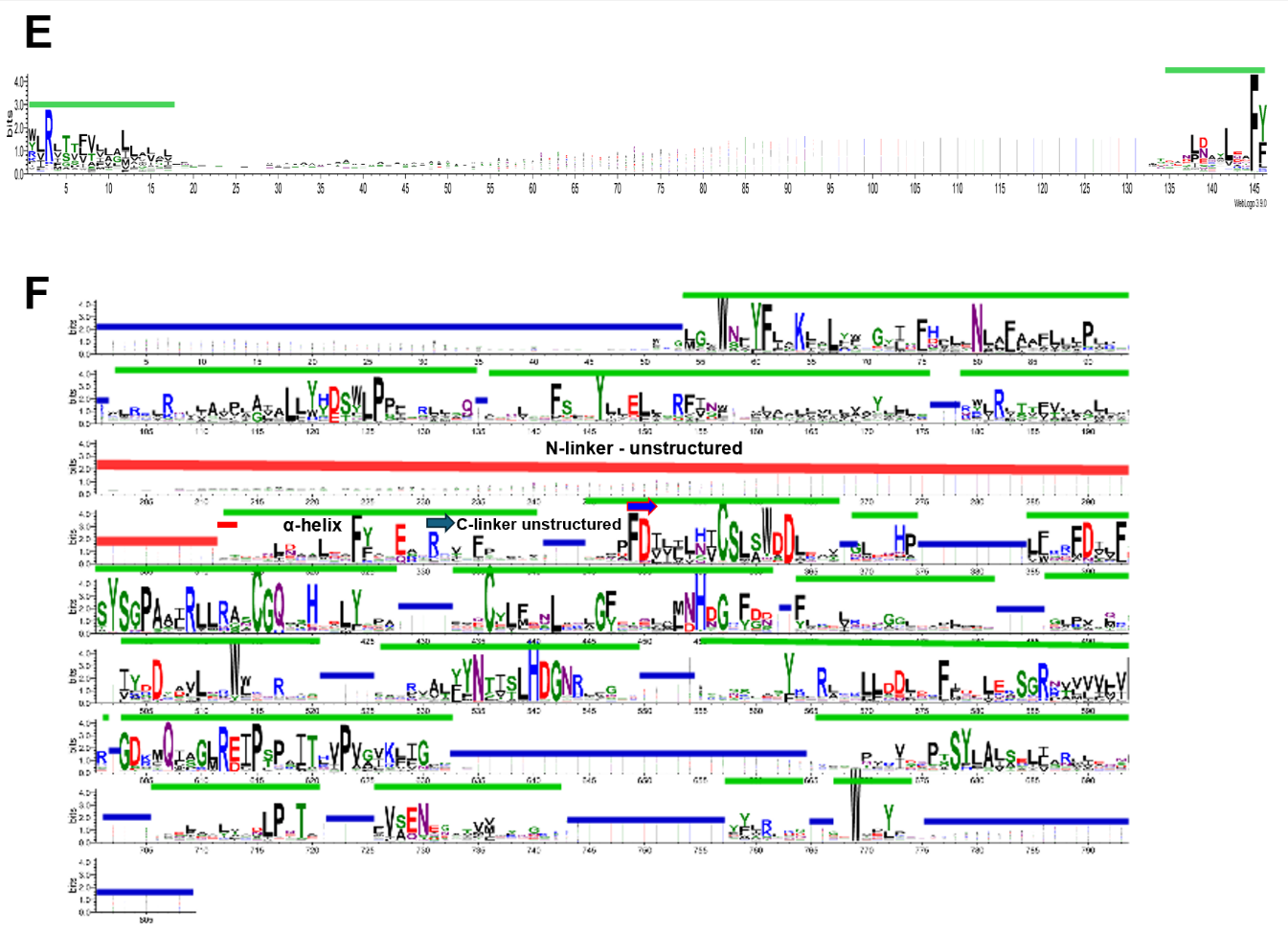

(**E**) Weblogo 3 of BcsG linker. Green line, amino acid sequences common to all BcsG proteins.

(**F**) Weblogo 3 of aligned BcsG proteins. Red bar, N-linker amino acid sequence; blue bar, region with amino acid sequences variable in sequence and length (including insertion observed only in a fraction of the proteins); green line, amino acid sequences common to all BcsG proteins.

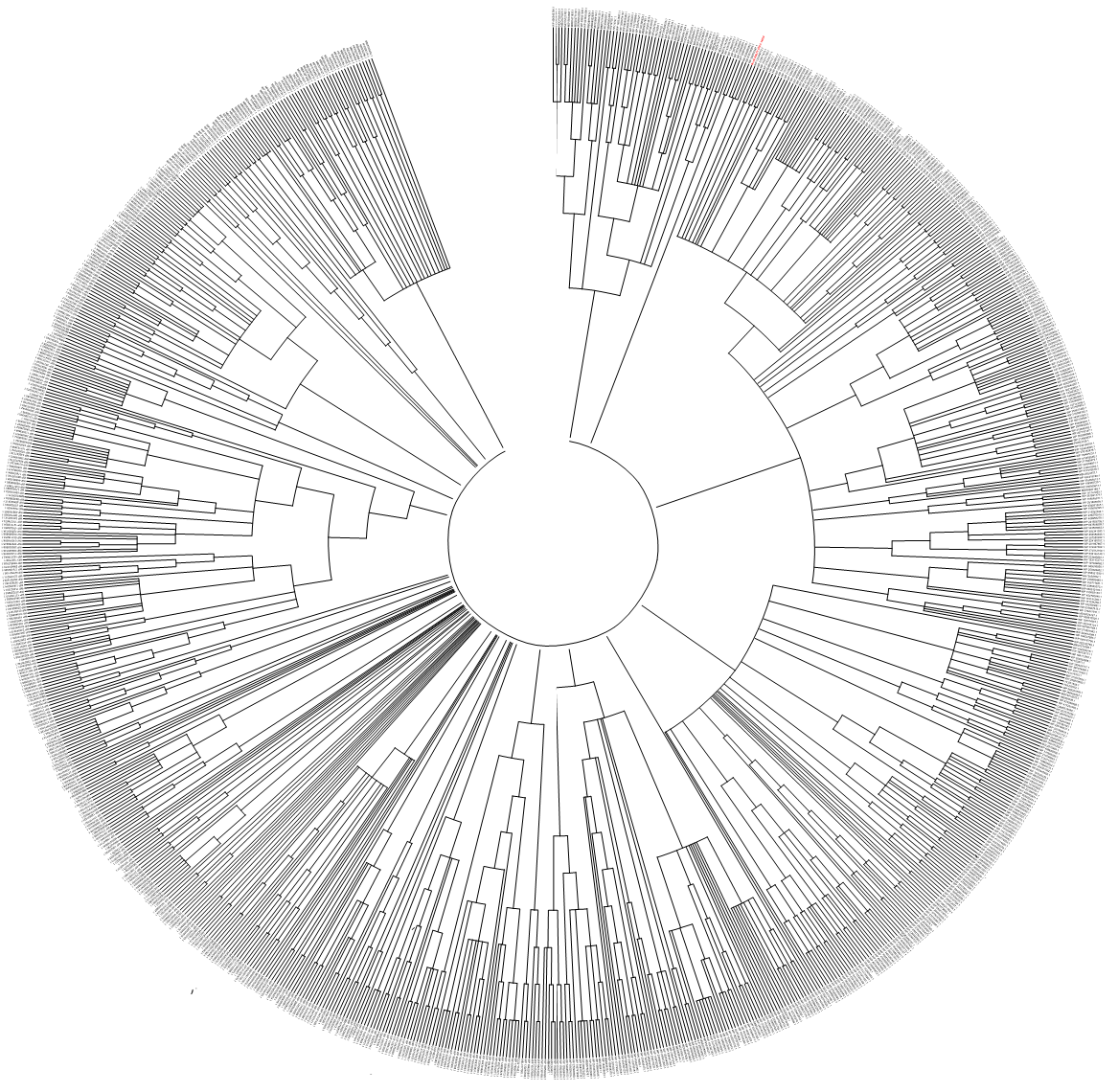

**Fig S3** Phylogenetic tree of BcsG homologs. BcsG homologs were retrieved from the NCBI database using BcsG of *S. typhimurium* ATCC 14028 as a query for Blast search (1). Sequences were aligned with ClustalX 2.1 (2) and manually curated in GeneDoc. A phylogenetic tree was constructed in MEGA 11.0 (3) with Maximum Likelihood with1000 bootstrap iterations. The resulting consensus tree was curated and displayed with InkScape.

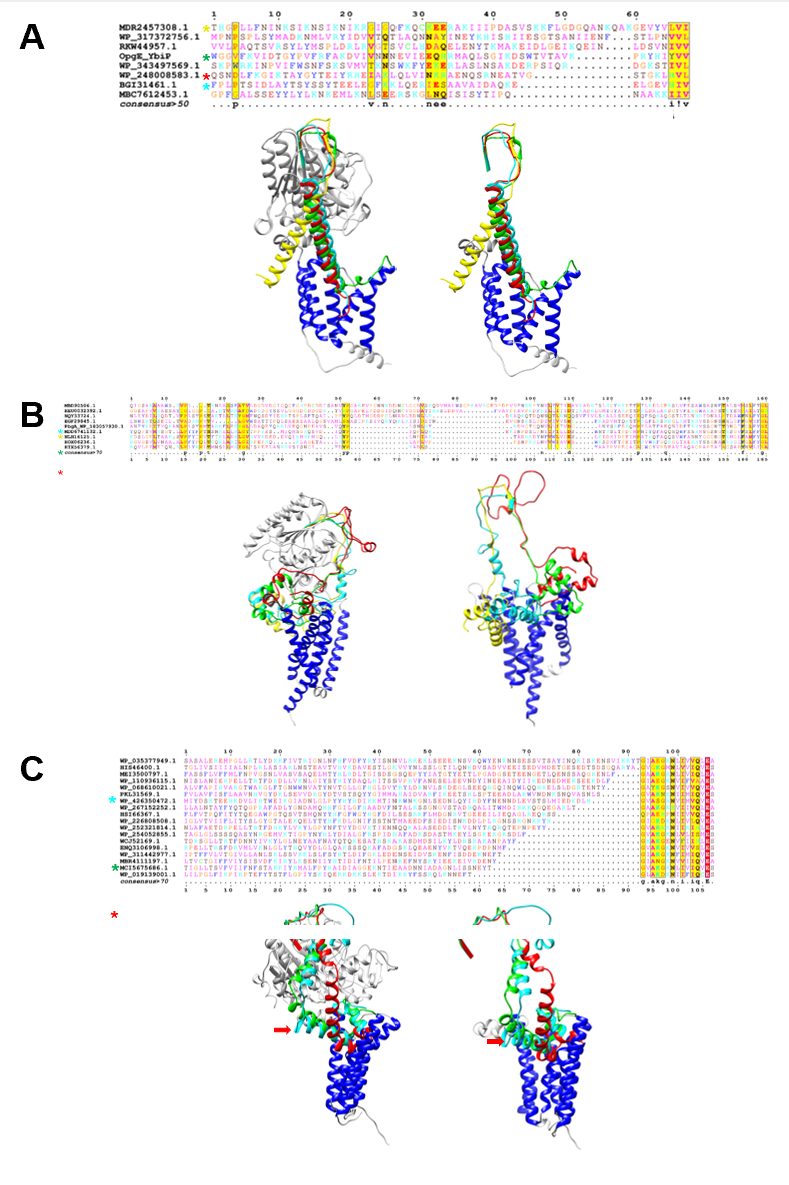

**Fig S4** Length span of fulllength linkers of homologs of alkaline phosphatase superfamily members OpgE, PbgA and LtaS. (**A**) Length span of linkers of homologs of alkaline phosphatase superfamily member OpgE and AlphaFold 3 models of OpgE of *S. typhimurium* ATCC 14028 compared to OpgE homologs MDR2457308.1 (yellow), WP_248008583.1 (red) and BGI31461.1 (cyan). Linker of OpgE in green. Only linker sequences are displayed for clarity for compared models. In dark blue, transmembrane helix of BcsG. (**B**) Length span of linkers of homologs of alkaline phosphatase superfamily member PbgA and AlphaFold 3 models of PbgA of *S. typhimurium* ATCC 14028 compared to PbgA homologs MBD90506.1, HEU0032392.1 and HIX56379.1. Only linker sequences are displayed for clarity for compared models. In dark blue, transmembrane helix of PbgA. (**C**) Length span of linkers of homologs of alkaline phosphatase superfamily member LtaS and AlphaFold 3 models of LtaS of *Staphylococcus aureus* PS47 compared to LtaS homologs WP_019139001.1 (red) and WP_03577949.1 (cyan). Linker of LtaS in green. Only linker sequences are displayed for clarity for compared models. In dark blue, transmembrane helix of LtaS.

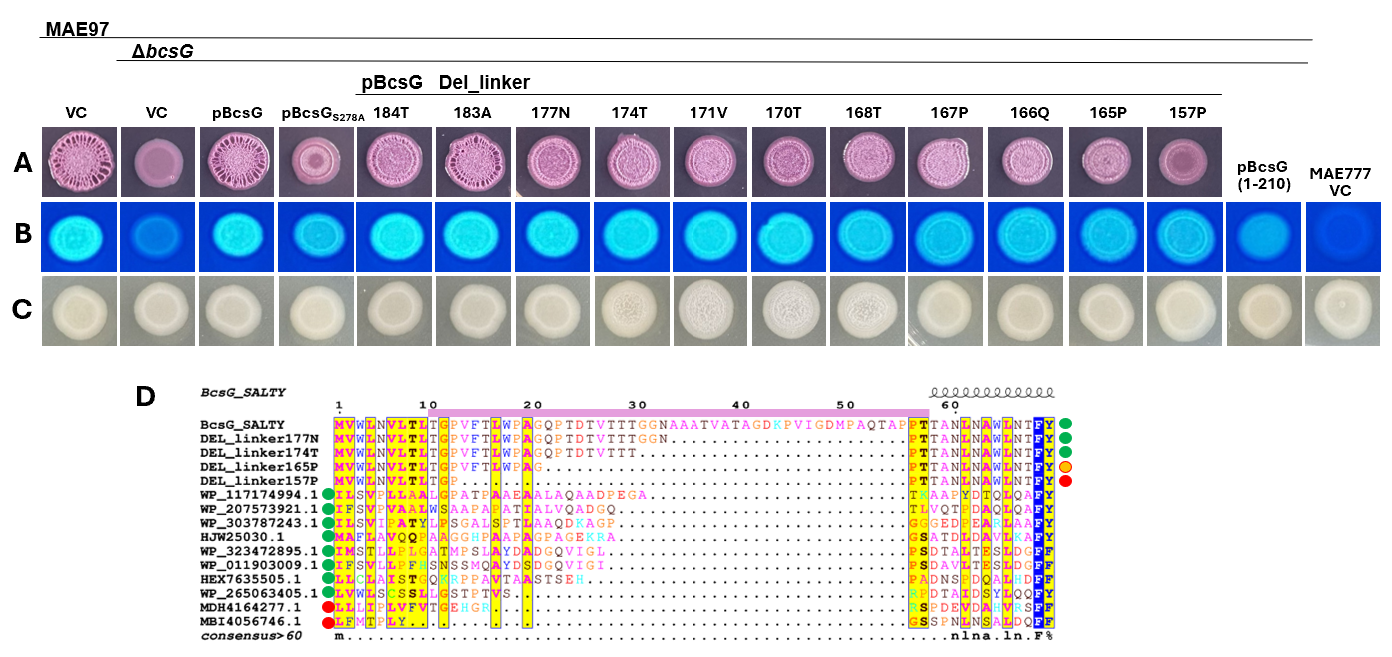

**Fig S5** Effect of BcsG N-linker length on colony morphology and in the phylogenetic context. (**A**) Congo red colony morphotype of *S. typhimurium* MAE97 Δ*bcsG* complemented by BcsG and its N-linker length variants*.* MAE97 vector control (VC) and MAE97 Δ*bcsG* pBcsG were positive, while MAE97 Δ*bcsG* VC and MAE97 Δ*bcsG* pBcsG S278A complemented with catalytically inactive BcsG were negative control references. Medium, LB without NaCl. VC=pBAD30. (**B**) Calcofluor white binding and (**C**) colony morphology of *S. typhimurium* MAE97 Δ*bcsG* complemented by BcsG and its N-linker length variants*.* MAE97 vector control (VC) and MAE97 Δ*bcsG* pBcsG were positive, while MAE97 Δ*bcsG* VC and MAE97 Δ*bcsG* pBcsG S278A complemented with catalytically inactive BcsG were negative control references. Medium, LB without NaCl. VC=pBAD30. (**D**) Comparison of N-linker variants of BcsG with BcsG homologs displaying the shortest N-linker sequences and co-localisation of BcsG homologs with cellulose biosynthesis gene clusters (+- five genes) indicated by green versus red circles on the left. Circles on the right side of the alignment: Observed phosphoethanolamine modification of the glucan chain by BcsG linker mutants. Red, no modification monitored; orange, residual modification monitored and green, substantial modification monitored. Light red bar above alignment, N-linker sequence of BcsG. Cells were grown on LB without salt agar plates at 37℃ for 24 h.

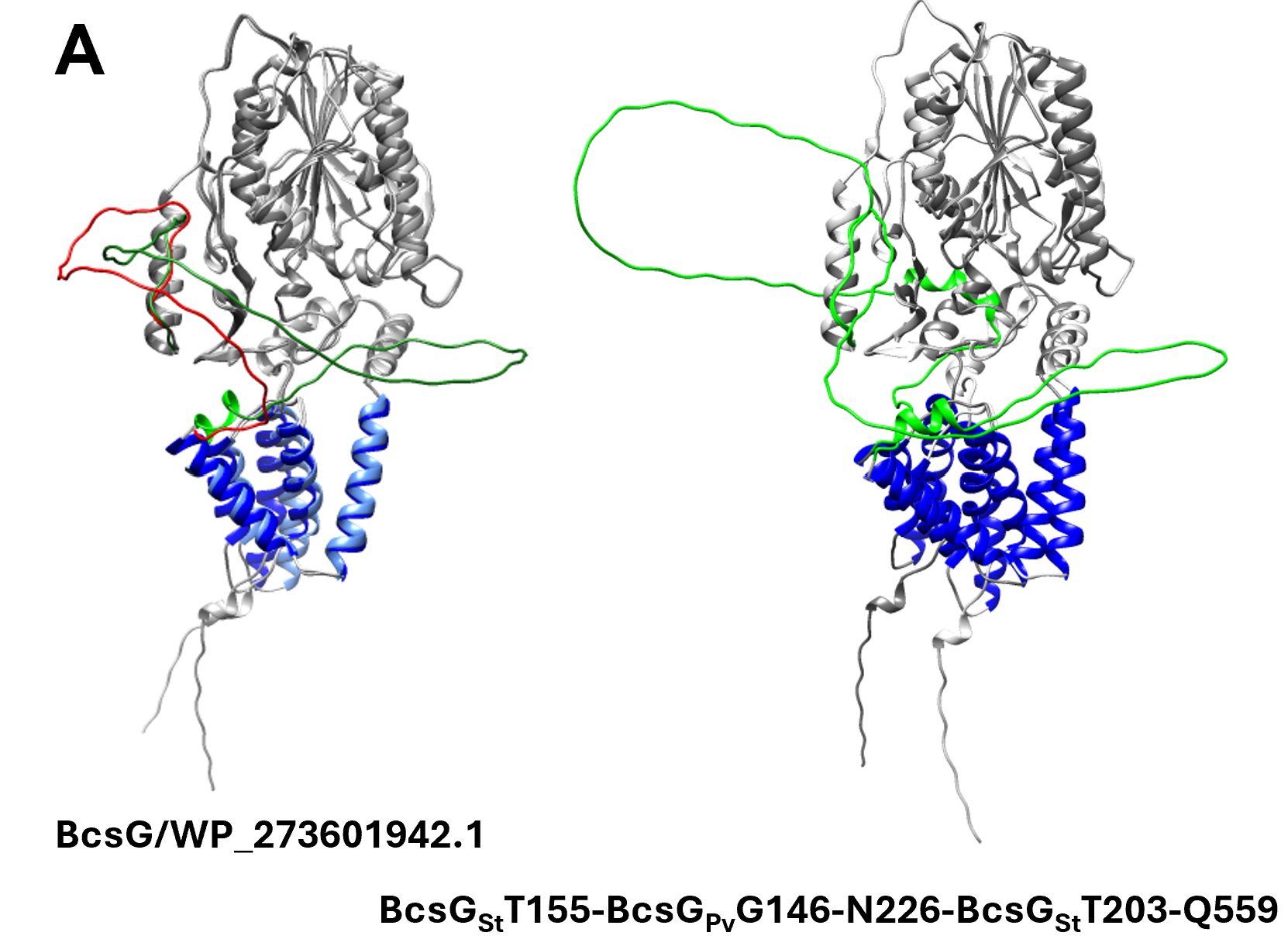

**Fig S6** Construction and AlphaFold 3 models of BcsG hybrid proteins. (**A**) Overlay of AlphaFold 3 models of BcsG_St_ from *S. typhimurium* ATCC 14028 with BcsG_Pv_ from *P. vulgaris* (WP_273601942.1). N-linker for BcsG_St_ in red and for BcsG_Pv_ WP_273601942.1 in green (left). Transmembrane helices in dark and light blue and periplasmic part in light and dark grey, respectively. Right, BcsG_St_ hybrid with BcsG_Pv_ (WP_273601942.1) linker G_149_GFTHFSQGMVPAVSANSFTQENSVNNEISSERASEIEVLPATVDSPGVTSMAPEVITKPVEQTVMGTLYPPQKHQFN_226_ in green replacing amino acids 156-202 of BcsG_St_. Two different predicted structures from two consecutive prediction events for BcsG_St_T155-BcsG_Pv_G146-N226-BcsG_St_T203-Q559 are overlayed. The structure of the N-linker sequences are predicted with predominantly very low per-atom confidence estimate of pIDDT<50.

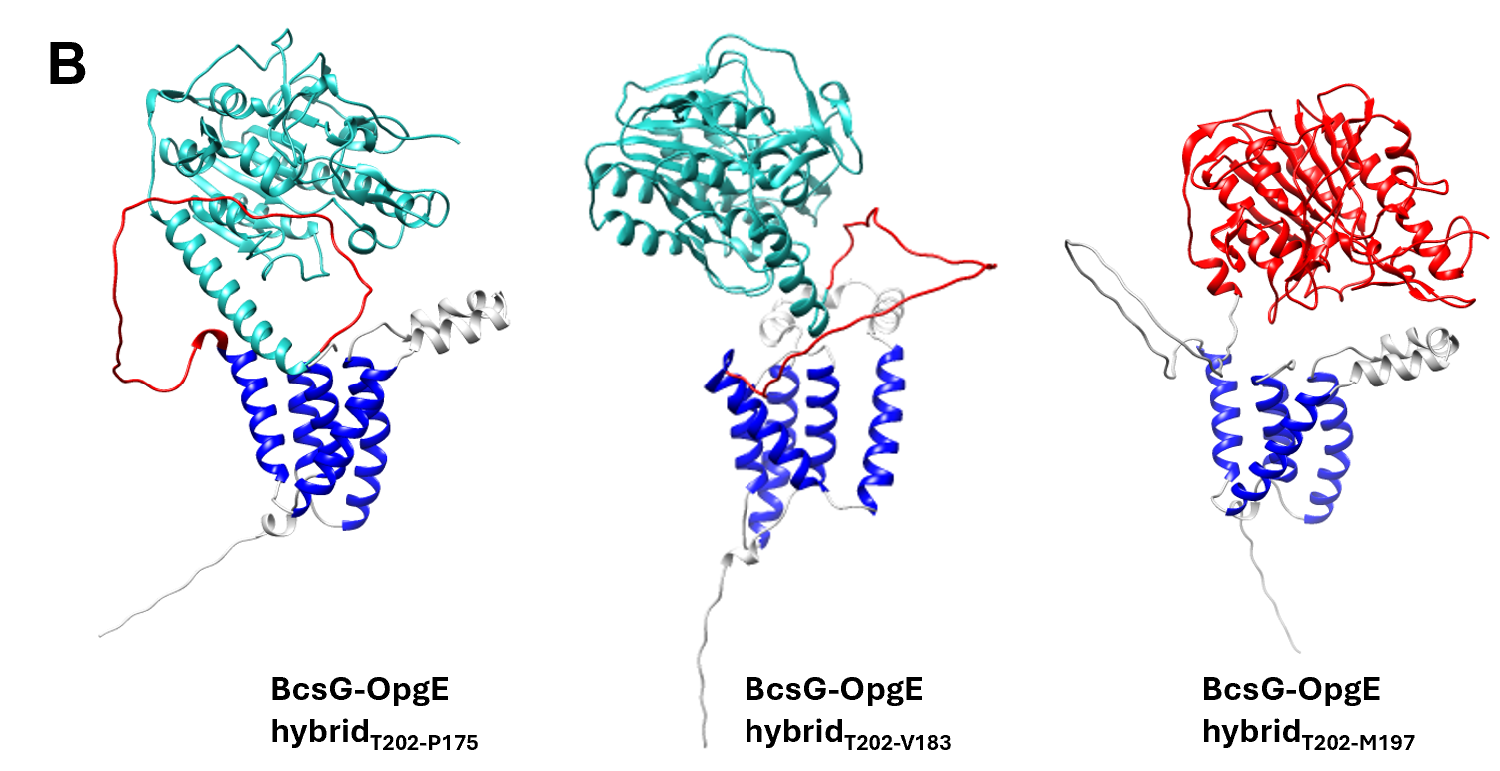

(**B**) AlphaFold 3 models of BcsG-OpgE hybrid protein with transmembrane part and N-linker from BcsG (in blue (transmembrane helices) and grey) and 3’ part of linker and catalytic domain from OpgE (in green or red).

**SUPPLEMENTARY MATERIAL**

**Codon optimized BcsG template from *Proteus vulgaris***

**GTACTGACATTAACC**

ggaggttttacccacttcagtcaggggatggttcccgcagtatctgccaattcctttactcaagaaaactccgttaataatgaaattagcagtgaacgggcctccgaaattgaagttcttcctgcgaccgtagattcgcctggcgttacgtcaatggcaccagaagttattaccaaaccagtcgaacaaacagttatgggcactttatat**c**ccccgcaaaaacatcagttcaacaacaaagtgttagacgattggcttactcaattttattcctatgaaaagaagcgtattactccatttcctgtacagctgtcagcagatgctcagccttttgatattctgattattaatatttgtagtttaagcacagctgatgccgcagcagttggtctgcaagaacaccctatatggggtaattttgatgtgttattcagtcatttcaacacagtgtcttcgtactcagggcctgcaagcctgcgtttattacgcgccagttgtggacagacccatcatagtgatctgtatgaccccacagatacacagtgccttttgatggataatttaagctctctgggatttgctaaaaaattagttctggatcataatggtaagtttgggaactatttgcaagaagtacaacaacttggcaatctgaatattgctctgcaagaccaagaaaatttaagccatcagattacagcctttgatggtactaaaatctataatgataaagaaaccttgatgcggtggcttcaaggccgtgaacagtcgaatgaaagtcggagtgtcacgttcgtgaatctggtatctttacacgacgggaaccggtttgtcggtgagaataacacagcggactatgggaaacgtgcctctacactgctggatagtttagataattttatgaacgaactggacaagaaaggcagaaaggtgatggtggttattgttcccgagcatggagccgctcttcttggtgataaaacgcaaatgagtggcctgagagatatcccgtcacagagtatcactacggtgcctgtcggtatccggtttacggggataaaagatcgcgcacagttctatccgccagtcattgtggaagaaccgagctcttacctggccttgtctgagttcattagccgcaatgtcaatggcgatgtatttaatcagtcaagtatcgattggaacgctttggcatctggcttaccgcaaacggcgaatgtggccgaaaaccaagccacgattgtggtggactatcaaggacaatcatacataaaactgaatggcggtgaatggatcaactatccgaac***CATCACCATCACCATCAT***taa**AAGCTT****GGCTGTTTTG**

in bold italics and upper case, codons coding for histidine;

in bold and upper case letter, nucleotide sequence homologous to pBAD28 vector

**Table 2.** Strains used in this study

| **Strain name** | **Genotype** | **Reference** |
| --- | --- | --- |
| ***Escherichia coli* strains** | | |
| TOP10 | F^-^ *mcrA* (*mrr-hsdRMS*-*mcrBC*) 80*lacZ* M15 *lacX74* *recA1 ara139* (*ara-leu*)*7697* *galU galK rpsL* (StrR) *endA1 nupG* | Invitrogen |
| DH5α | F– φ80 lacZΔM15 Δ(lacZYA–argF)U169 recA1 endA1 hsdR17(rK–, mK+) phoA supE44 λ– thi-1 gyrA96 relA1 | New England Biolabs |
| ***S. typhimurium* ATCC 14028 derivatives** | | |
| UMR1 | ATCC 14028s rdar_28°C_, Nal^r^ | (4) |
| MAE97 | UMR1 p*csgD*1 Δ*csgBA*102, pEtN-cellulose (pdar) 28/37 °C | **(5)** |
| MAE14 | UMR1 Δ*csgBA*101::Km^r^, pEtN-cellulose (pdar) 28°C | **(5)** |
| MAE97Δ*bcsG* | UMR1 p*csgD*1 Δ*csgBA102*, Δ*bcsG* | (6) |
| MAE1264 | MAE97 *bcsA*-3xFLAG | (7) |
| MAE1264 Δ*bcsG* | MAE97 *bcsA*-3xFlag, Δ*bcsG* | (7) |
| MAE777 | UMR1 p*csgD*1 Δ*csgBA*102, *ΔbcsA102* p*csgBA::*Km | (8) |

**Table 3.** Primers used in this study

| **Name** | **Sequence** | **Purpose** |
| --- | --- | --- |
| BcsG_del linker 157P_200P_F | CCGACGACCGCGAATCTGAAC | Construct Del_linker 157P |
| BcsG_del linker 157P_200P_R | CGGGCCGGTTAATGTCAGTAC |  |
| BcsG_del linker 165P_200P_R | GCCTGCCGGCCACAGCGTAAAAAC | Construct Del_linker 165P |
| BcsG_del linker 167P_200P_R | TGGCTGGCCTGCCGGCCACAG | Construct Del_linker 167P |
| BcsG_del linker 168T_200P_R | GGTTGGCTGGCCTGCCGGCCAC | Construct Del_linker 168T |
| BcsG_del linker 174T_200P_R | AGTTGTCGTCACCGTATCGGTT | Construct Del_linker 174T (170T) |
| BcsG_del linker 171V_200P_R | CACCGTATCGGTTGGCTGGCCTG | Construct Del_linker 171V |
| BcsG_del linker 177N_200P_R | GTTACCGCCAGTTGTCGTCACC | Construct Del_linker 177N |
| BcsG_del linker 183A_200P_R | CGCGACGGTAGCGGCC | Construct Del_linker 183A |
| BcsG_del linker 184T_200P_F | ACAGCGGGCGATAAGCCGG | Construct Del_linker 184T |
| Δlinker166Q-F | GTGGCCGGCAGGCCAG ccgacgaccgcgaatctgaac | Construct Del_linker 166Q |
| PCR-OpgE-F | ATGAATTCAACCCTTATCGATTC | Amplify *opgE* |
| PCR-OpgE-R | TTATTTTAGTGAAAAAATATCGCTGC |  |
| Vector-F_v2 | ACTGGGTGCCTTACCCGCAG | Amplify linearized plasmid pBAD30 |
| Vector-R | ATAGAAGGTGTTCAACCAGGCG |  |
| 197M-F-Insert | CCTGGTTGAACACCTTCTATatggcgcagctatccgg | Construct *bcsG*-hybrid 197M (opgE) |
| R_insert | AAACAGCCAAGCTTTTAATGATGATGATGATGATGttttagtgaaaaaatatcgctgccc |  |
| 175P-F | CCTGGTTGAACACCTTCTATccggtttttcgcttcgccaaag | Construct *bcsG*-hybrid 175P (opgE) |
| 184I-F | CCTGGTTGAACACCTTCTATatcgtcaataacaacgaagtcatt | Construct *bcsG*-hybrid 184I (*opgE*) |
| Linker-l-f | TGAATGTACTGACATTAACCggaggttttacccacttcagtca | Construct *bcsG*-hybrid-l-linker (*Proteus vulgaris*) |
| Hybrid bcsG control 1-F | CCATGGGCGCTTTCGGCG | PCR confirmation primers of *bcsG*-hybrid (*opgE*) |
| New hybrid gene-R | CGCCGCTATGACGAAGCTGAGTATAGAG |  |

Homology regions for in vivo cloning are shown in lowercase, and template-complementary sequences are shown in uppercase.

**Table 4.** Plasmids used in this study

| **Plasmid name** | **Features** | **Reference/Source** |
| --- | --- | --- |
| **Empty plasmids** | | |
| pBAD30 | pACYC184 origin, M13, Amp^r^, P_BAD_ promotor, L-arabinose-inducible | **(9)** |
| **Template plasmid** | | |
| pBAD-BcsG | *bcsG* cloned in XbaI/HindIII sites in pBAD30 with a 6x His Tag | (6) |
| **New plasmid constructs** | | |
| pBAD30-bcsG Del_linker 184T | Linker deletion | This study |
| pBAD30-bcsG Del_linker 183A | Linker deletion | This study |
| pBAD30-bcsG Del_linker 177N | Linker deletion | This study |
| pBAD30-bcsG Del_linker 174T | Linker deletion | This study |
| pBAD30-bcsG Del_linker 171V | Linker deletion | This study |
| pBAD30-bcsG Del_linker 168T | Linker deletion | This study |
| pBAD30-bcsG Del_linker 167P | Linker deletion | This study |
| pBAD30-bcsG Del_linker 166Q | Linker deletion | This study |
| pBAD30-bcsG Del_linker 165P | Linker deletion | This study |
| pBAD30-bcsG Del_linker 157P | Linker deletion | This study |
| pBAD30-*pssA* | *pssA* gene inserted in XbaI and SphI sites, C‑terminal 6xHis Tag | This study |
| pBAD30-*pgsA* | *pgsA* gene inserted in XbaI and SphI sites, C‑terminal 6xHis Tag | This study |
| pBAD30-bcsG hybrid 197M | OpgE (aa 197-214) replacing BcsG (aa 156-202) | This study |
| pBAD30-BcsG hybrid 184I | OpgE (aa 184-214) replacing BcsG (aa 156-202) | This study |
| pBAD30-BcsG hybrid 175P | OpgE (aa 175-214) replacing BcsG (aa 156-202) | This study |
| pBAD30-BcsG hybrid l-linker | BcsG_Pv_ linker from *P. vulgaris* (aa 249-224) replacing BcsG (aa 156-202) | This study |
